# Rankings of tuberculosis antibiotic treatment regimens are sensitive to spatial scale, detection limit, and initial host bacterial burden

**DOI:** 10.1101/2025.04.10.648004

**Authors:** Christian T. Michael, Maral Budak, Pauline Maiello, Kara Kracinovsky, Mark Rodgers, Jaime Tomko, Philana Ling Lin, JoAnne Flynn, Jennifer J. Linderman, Denise Kirschner

## Abstract

Pulmonary infection after inhalation of Mycobacterium tuberculosis (Mtb) causes tuberculosis (TB). TB presents with lung granulomas - complex spheroidal structures composed of immune cells and bacteria. Granulomas often have centralized caseum (necrotic tissue) where mycobacteria are quarantined, complicating and prolonging multi-antibiotic regimens. Determining which antibiotic regimens are optimal for reducing treatment time and toxicity is a goal of recent TB eradication campaigns. Clinical trials are expensive and challenging, making it difficult to untangle which host-pathogen interactions drive the heterogeneous infection and treatment outcomes observed at between-host and within-host scales. To determine responses to antibiotic regimens, we simulate treatments in *HostSim*, our whole-host mechanistic, multi-scale computational model of Mtb-infection. *HostSim* tracks dynamics of pulmonary Mtb-infection over molecular, cellular, tissue, organ, and whole-host scales. We create a heterogenous virtual cohort, comprising distinct hosts, for virtual clinical trials. We represent drug treatments by newly-integrating pharmacokinetics / pharmacodynamics into HostSim, simulating treatment with commonly-prescribed TB antibiotic regimens (e.g., HRZE or BPaL). Our approach allows us to identify both (1) which hosts/granulomas most improve with treatment, and (2) which mechanisms influence outcome heterogeneity. By tracking experimental and clinical measurements, we virtually recreate several drug rankings from literature. We find that many methods of ranking treatment efficacy are strongly influenced by the ‘definition of improvement’ used and, in some cases, the detection threshold of CFU. Other rankings depend on initial bacterial burden of hosts/granulomas. Our work suggests that metrics for regimen optimality may be orthogonal, which could explain seemingly-contradictory findings from prior studies.

## 1. Introduction

Tuberculosis (TB) is both an ancient disease and the current leading cause of death by a single infectious disease in the world, causing 1.25 million deaths in 2023 alone^1,2^. TB is caused by the inhalation of the bacillus *Mycobacterium tuberculosis* (Mtb) that mainly infects lungs and lymph nodes and but can include other clinical manifestations, including various extrapulmonary organ involvement (e.g., liver, brain, kidney, spine). Pulmonary Mtb infection leads to the formation of multiple lung granulomas, hallmark structures arising from infection with Mtb that are composed of host immune cells, bacteria, and dead cellular debris. Because of spatial quarantining of Mtb within granulomas, approximately 90% of Mtb-infected humans have subclinical Mtb infection: a chronic, asymptomatic disease state^3,4^ that reactivates in ∼10% of individuals^5^. Granulomas are diverse microenvironments within which Mtb adapts, generating a range of heterogeneous Mtb phenotypes^6^. To target various Mtb phenotypes simultaneously and to reduce the risk of development of resistant bacteria, TB treatment involves administration of multiple antibiotics simultaneously^7^. The standard treatment for drug-susceptible Mtb infection is the administration of four drugs – isoniazid (H; INH), rifampicin (R; RIF), pyrazinamide (Z; PZA), and ethambutol (E; EMB) – over 6-9 months^8^.

The World Health Organization’s (WHO) EndTB strategy aims to reduce the TB death toll by 95% by 2035^9^. To achieve this goal, many research efforts focus on discovering or repurposing antibiotics that may increase the cure rate. Due to an increased number of anti-Mtb antibiotics and the need for combination therapy, this leads to a large number of possible TB regimens to be tested, rising to 10^17 10^. To overcome feasibility challenges and screen regimen combinations efficiently, we developed an *in silico* multi-scale mechanistic model (*GranSim*) that simulates host-immune responses upon Mtb infection in lung tissue and granuloma formation as a result of interplay between host immune response and mycobacteria^11-15^. We have continuously curated *GranSim* to data generated from NHPs, which have similar immunological dynamics as humans during Mtb infection and granuloma formation^16-20^. Moreover, we incorporated a pharmacokinetics (PK) and pharmacodynamics (PD) model of anti-Mtb antibiotics within *GranSim* and simulated multiple regimens with different drug combinations^21-23^. Treatment outcomes simulated in *GranSim* reflect both human clinical trials as well as *in vivo* studies supporting credibility of our model^24,25^. *GranSim* is a multi-scale model that spans molecular to tissue scales; however, it lacks host-scale outcomes of Mtb infection, such as the dynamics of multiple granulomas and granuloma dissemination, which can significantly affect treatment. Due to these limitations, it is challenging to predict host-scale treatment outcomes using *GranSim*.

We recently developed a whole host-scale, mechanistic *in silico* model of TB disease, *HostSim*^26,27^. *HostSim* captures pulmonary TB dynamics within 3 compartments: lungs, blood and lung-draining lymph nodes. It represents multiple granulomas within a single host, capturing dynamics within both the cellular regions of granulomas and their caseous necrotic cores. We calibrate this model using NHP datasets as well as synthetic data generated from *GranSim*. Here we adapt our PK/PD modeling of antibiotics, including drug-drug interactions, in *GranSim* for incorporation within *HostSim*, including the eight most prescribed antibiotics for TB: INH, RIF, PZA, EMB, Bedaquiline (B; BDQ), pretomanid (Pa; PTM), linezolid (L; LZD), and moxifloxacin (M; MXF). Unique to our framework is the ability to observe drug action and effects at molecular (drugs, cytokines, etc.), cellular (immune cells and bacteria), tissue (granulomas), host (lungs, lymph nodes and blood), and population (virtual cohort) scales.

We validate our *HostSim* treatment simulation framework in discrete steps. First, we show that *HostSim* captures Mtb infection features and dynamics from human and NHP data before treatment. Second, we calibrate our *HostSim* PK model to experimental PK data from literature for each of eight drugs. Third, we compare *HostSim* output to bacterial burden from NHPs treated with three well-studied regimens: HRZE, RMZE, and BPaL. Fourth, we perform a multi-scale sensitivity analysis that identifies drivers of host heterogeneous treatment responses in the same three regimens, and these reflect key mechanisms identified in literature. Finally, we recapitulate rankings for several drug regimens that have been previously established using clinical data, NHP data, and granuloma-scale simulation data.

In this work, our goal is to create a virtual cohort that can be used for testing a number of different antibiotic regimens and then for ranking those regimens. To this end, we define a *virtual cohort* as a collection of virtual Mtb-infected hosts that we can study *in silico* under multiple scenarios, and we define *virtual clinical trials* as simulations of the virtual cohort undergoing treatment to characterize infection outcomes while remaining accurate to available human and animal data. We also highlight the orthogonality of different ranking methods of drug efficacy, even when these rankings superficially and intuitively may be measuring the same clinical feature—e.g., sterilizing potential. To create multiple rankings, we use data from a single virtual cohort that we treat with three sets of antibiotic regimens, studied in previous work, to create a single underlying “ground truth” from which to rank drug efficacy. In these simulations, we repeat data collection methods from previous experimental studies or clinical trials. We find that in some cases, using different methods to rank CFU sterilizing potential has poorly correlated outcomes. In one ranking analysis, varying the limit of detection of bacteria between 1 CFU per granuloma and 10 CFU per granuloma substantially changes relative rankings among twelve drug regimens. At the host scale, we find that ranking only high-CFU-burden hosts (or low-CFU-burden hosts, mimicking different clinical trial admission criteria) results in anti-correlated rankings. Ultimately, we demonstrate that subtle assumptions in experimental design may result in apparent contradictions if one assumes that ranking methodologies are interchangeable, rather than contextual.

## 2. Results

### 2.1 *HostSim* captures the spectrum of TB progression dynamics within a virtual cohort

As a validation of our *HostSim* model and as a foundation for our virtual clinical trials, we seek to ensure that *HostSim* reproduces dynamics of both subclinical and active pulmonary Mtb infection without treatment. After calibrating *HostSim* using CaliPro with NHP datasets (see Methods), we generate a virtual cohort of 500 virtual hosts. We seed each virtual host with 13 granulomas at time and simulate disease progression until 480 days post-infection (p.i.) (**Figure 1**). By one-year p.i., we find that 12.2% of our virtual cohort is actively infected, 86.2% exhibit subclinical infection, and 1.2% are apparently-sterilizing (See Methods for virtual host classification). This is similar to ranges observed in humans, which have ∼90% subclinical infection and ∼10% active disease, with very few fully sterilizing patients^3,4^, although determining the precise relation between true sterilization and latent disease is extremely difficult^28^. Without treatment, only 1 out of 8 hosts that are apparently-sterilizing are actually *sterilized*; the other 7 virtual hosts have fewer than 20 Mtb trapped within caseum. This indicates that while these granulomas are highly successful in controlling infection, the hosts may be at risk of long-term reactivation (in the event that they do in fact harbor live Mtb^28^). We notice that in most subclinical virtual host cases, intracellular CFU counts stabilize below approximately 100, and a larger reservoir of CFU are trapped in caseum. By contrast, virtual hosts with active disease have high intracellular Mtb levels compared to hosts with subclinical infection (**Figure 1**B). In hosts with active disease, we find higher levels of intracellular Mtb populations and uncontrolled extracellular growth outside of caseum. This recapitulates the common understanding that Mtb-controlling non-sterile granulomas trap Mtb within caseum; indeed, recent studies show that dead macrophages in normoxic environments provide an ideal replicative niche to Mtb^29^.

**Figure 1.**
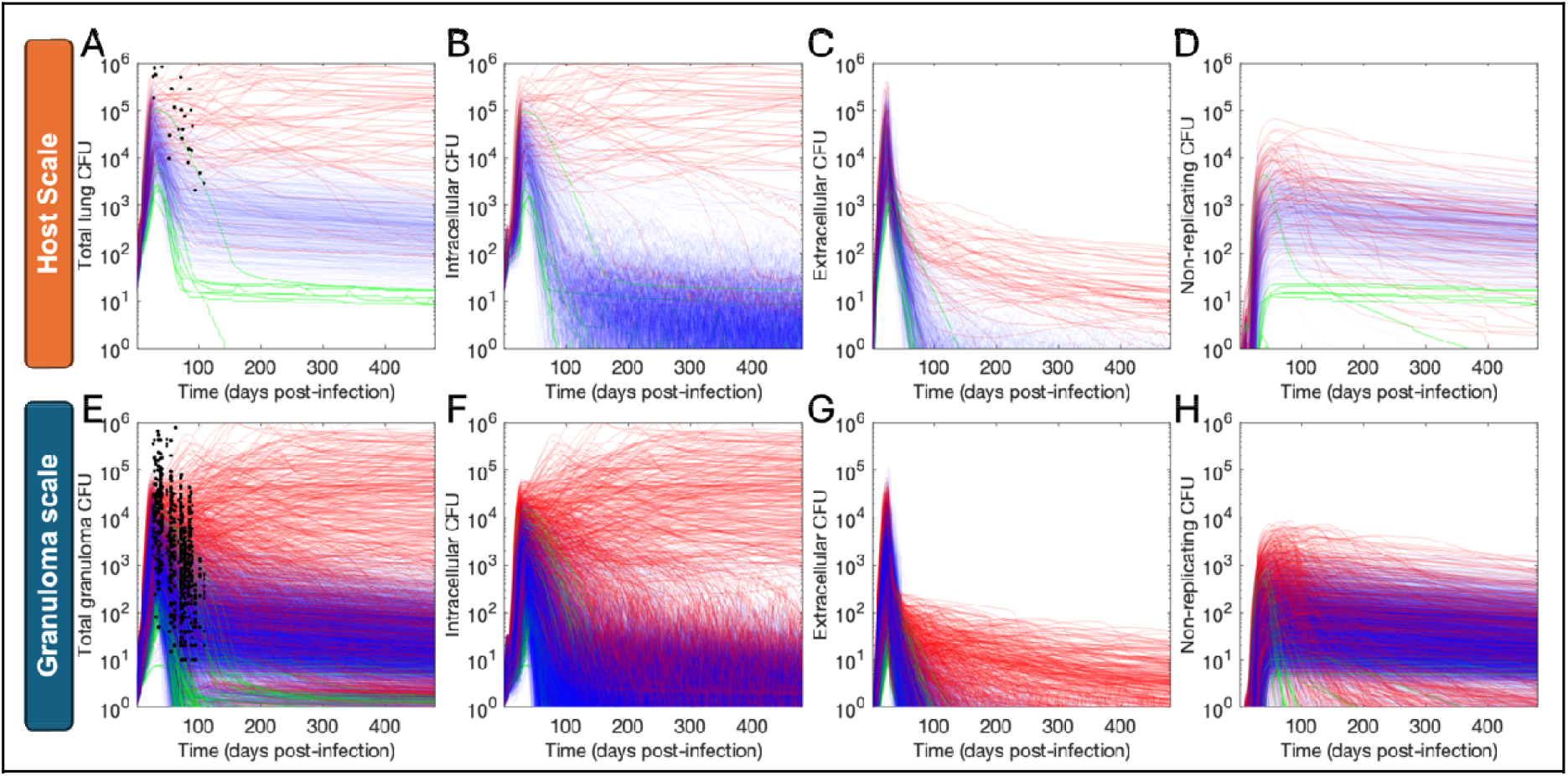
Multi-scale bacterial burden within 500 virtual hosts. Log-scale plots of CFU from virtual hosts (A-D) and their corresponding primary granulomas () (E-H). Each curve represents a CFU trajectory over 480 days post-infection for an individual host or granuloma. In each column we show a different heterogenous subpopulation of Mtb by spatial location: total bacterial burden (A, E), intracellular CFU (B,F), extracellular CFU (C,G), and non-replicating CFU trapped within caseum (D,H). All lines are colored based on the clinical classification of the virtual host: apparently-sterilizing (green), subclinical Mtb infection (blue), and active TB (red). (A, E) For validation, CFU counts measured from NHP lungs and granulomas in previous experimental studies are also shown (black dots in A, E)^30-33^.

### 2.2 *HostSim* virtual dosing captures the pharmacokinetics of multiple anti-TB antibiotics

To perform virtual drug studies we include both PK and PD sub-models within *HostSim* (see Methods) as we have done previously for our granuloma scale model, *GranSim*^21-25,34^. We then use datasets derived from rabbit^35-37^ and NHP models^38^ (both generate human-like, caseous granulomas); as well as data from human samples^39^ to calibrate our PK model for 8 drugs: for INH, RIF, PZA, EMB, BPA, PTM, LZD, and MXF. We administer each drug to the same virtual cohort of 500 hosts (and consequently the same 6500 primary granulomas) using our multi-scale PK model (see Methods for dosing and other details). As these PK measurements come from a variety of dosages and times post-dosing, we present a summary of our calibration. Our model predicts drug concentrations that fall at least within one order of magnitude of experimentally observed drug concentrations and much closer in most cases (**Figure 2**). Our ability to match PK against a wide range of data from human and animal samples gives us confidence in utilizing the model for virtual clinical trials.

**Figure 2.**
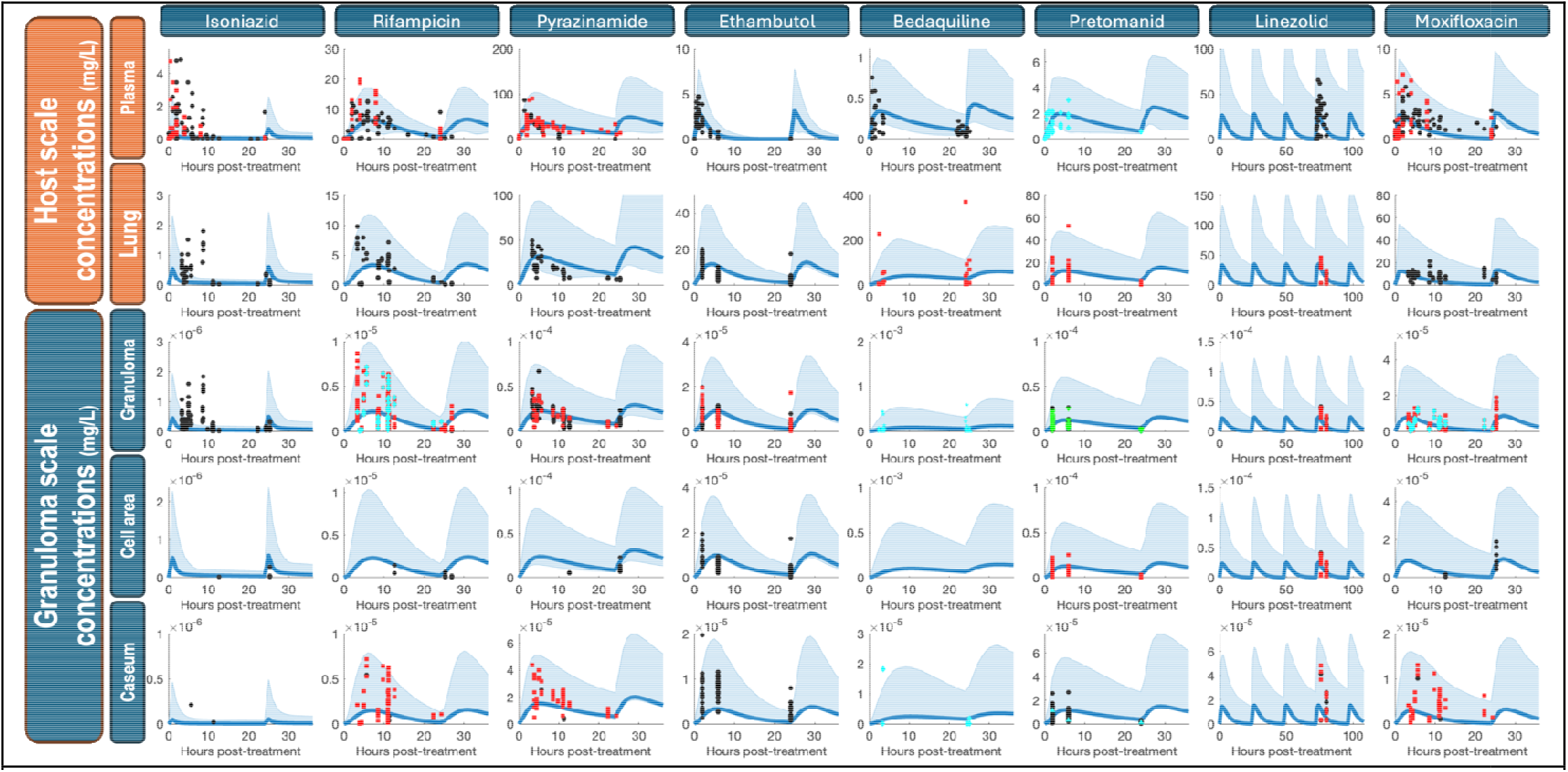
Multi-scale calibration of antibiotic pharmacokinetics using 500 virtual hosts. PK calibrations for multiple antibiotics across different physiological compartments are tracked within *HostSim*. Our linear-scale ‘ribbon plots’ depict the median (bold line), 1st and 99th percentile (blue ribbon) for drug concentrations from virtual hosts (rows with orange heading) or granulomas (rows with blue heading). Plots in each column and row summarize the distribution of concentrations for each of the drugs and compartments, respectively. Each point on the superimposed scatterplots indicates experimental data directly comparable to our simulation; different symbols and colors within a plot indicate that data came from different studies from literature. We use rabbit, NHP, and human PK datasets derived from literature for INH, RIF, PZA, EMB, BDQ, PTM, LZD, and MXF^35-37,39^.

Pharmacokinetics are only the first step in modeling drug dynamics within a host. It is also necessary to capture the action of the drugs through pharmacodynamics. Our PD model uses the concentrations from the PK predictions to calculate killing rates that are then implemented in the three subpopulations (i.e., bacteria in each spatial regions of the granuloma); these calculations for killing rates are identical to those we have used previously^22,40^ (see Methods). We recently calibrated our PD model using data from bactericidal assays wherein Mtb is exposed to drugs in different physiological conditions^24,25^ (e.g., caseum homogenate, macrophage assays, and standard media; see Methods for details). We apply and validate our combined PK/PD implementation in the following sections.

### 2.3 *HostSim* virtual clinical trials recapitulate the dynamics of HRZE, BPaL and RMZE drug regimens

With our PK and PD models in place, we next validate our model of drug treatment, showing predictions are consistent with multiple existing datasets. We examine the regimens HRZE, BPaL, and RMZE, as these regimens have been extensively studied (see Methods for dosing details)^41-49^. First, we observe sterilization curves (**Figure 3**) that qualitatively agree with previous observations of HRZE, BPaL, and RMZE^25,40-42,45,48,49^. In particular, we predict that (a) HRZE is the slowest of the three regimens to sterilize infections, (b) RMZE sterilizes some hosts very quickly, leading to a “plateau” sterilization profile, and (c) BPaL takes time to sterilize hosts but eventually reaches higher levels of sterilization than HRZE or RMZE.

**Figure 3.**
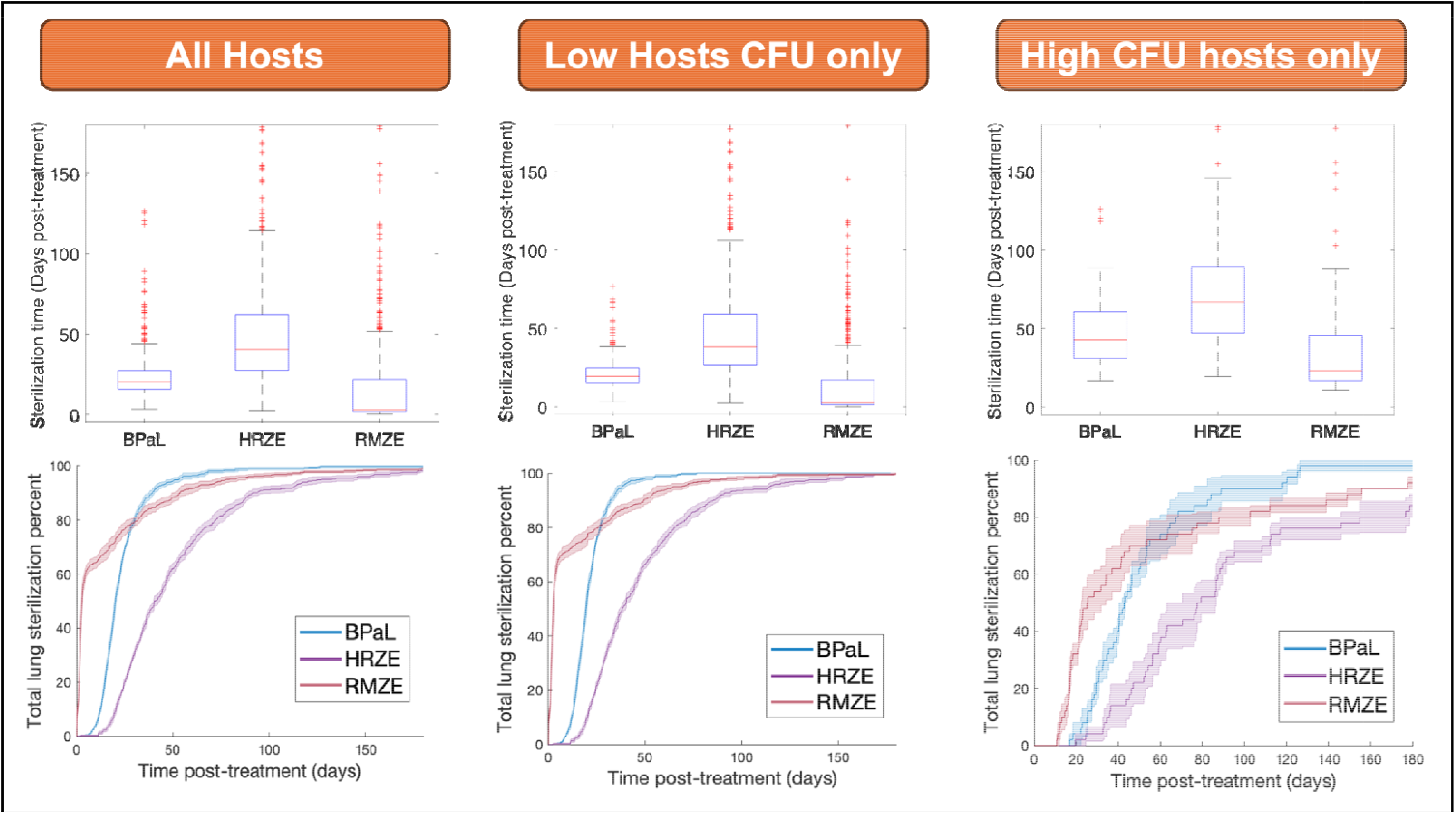
Efficacy of three drug regimens in virtual trials, as observed at the population scale. We test three regimens, RMZE, BPaL, and HRZE. We plot the number of hosts sterilized by (**left column**) all 496 hosts that remain infected by the start of treatment (300 days p.i.), (**middle column**) 443 low-CFU hosts (<10,000 total Mtb prior to treatment), and (**right column**) 53 high-CFU hosts (>10,000 total Mtb prior to treatment). (**Top row**) Distributions of times-to-sterilization by regimen measured in days p.i. (**Bottom row**) Ribbon curves show mean and standard error of sterilization time by regimen. We measure this by randomly dividing the virtual population into 5 equally-sized subgroups (each group n=99 for all-hosts analysis, n=86 low-CFU host analysis, n=11 high-CFU host analysis) and report the mean and standard error of sterilization percents per-subgroup over time (see Methods).

We validate our model at the population scale by comparing simulated drug efficacy predicted by *HostSim* with experimental and clinical Early Bactericidal Assays (EBAs)—a measure of bacterial reduction over time that is commonly reported^50^. We report this with two caveats: first is that clinical EBAs are measured with sputum and our virtual EBAs are measurements of whole-lung CFU; we assume that these measures are directly proportional in our comparison. Secondly, we calculated virtual EBAs both including and not-including non-replicating bacteria () (see Methods for virtual EBA calculation). As in the meta-analysis by Bonnett et al.^50^, we simulate EBAs at 2-, 7-, and 14-days post-treatment (**Table 1**). For HRZE, our mean and median EBA values including fall within the respective 95% confidence intervals reported by Bonnett at 2, 7, and 14 days, demonstrating a good match between our model and clinical findings. Interestingly, the confidence interval we compared to for EBA 0-14 is [0.1, 0.21]^50^, and our simulated EBA without is near the bottom of this range while without including is near the top of this range. This may indicate that a fraction of non-replicating bacteria consistently contributes to EBA measurements. Moreover, our simulated EBA values from BPaL are similar to that of HRZE, as observed in clinical trials^51^, which further validates our simulated antibiotic regimen efficacies. We note that RMZE had higher EBA than either BPaL or HRZE, especially during early infection. This is consistent with high levels of bactericidal activity seen in MXF^39,41^ which, during early infection, will be highlighted as EBA is more impacted by quickly killing large numbers of easily-accessible Mtb. These results demonstrate that *HostSim* well-captures bactericidal activity during early infection.

**Table 1:** Simulated EBA values for three antibiotic regimens. In plain text, we report EBA values computed assuming that total lung CFU (including non-replicating bacteria) contributes to the EBA measurement. In brackets and in bold, we include mean and median simulated EBA values computed assuming that non-replicating bacteria do not contribute to EBA.

| Regimen | EBA 0-2 (Mean, Median)<br>(With ) [Without ] | EBA 0-7 (Mean, Median)<br>(With ) [Without ] | EBA 0-14 (Mean, Median)<br>(With ) [Without ] |
| --- | --- | --- | --- |
| HRZE | (0.18, 0.13) [0.35, 0.37] | (0.13, 0.11) [0.26, 0.27] | (0.11, 0.10) [0.2, 0.21] |
| BPaL | (0.20, 0.18) [0.29, 0.30] | (0.16, 0.15) [0.21, 0.22] | (0.15, 0.14) [0.18, 0.18] |
| RMZE | (0.68, 0.43) [0.37, 0.37] | (0.32, 0.34) [0.28, 0.28] | (0.20, 0.22) [0.22, 0.23] |

### 2.4 *HostSim* virtual clinical trials show a multi-phasic decline of host and granuloma CFU that captures non-human primate data

We next validate outcomes of simulated treatment in *HostSim* against datasets from *Cynomolgus macaques*, which provide detailed and human-like data at both the granuloma and whole-host scales^52^. Specifically, we compare virtual to NHP CFU counts for three well-studied regimens: HRZE, RMZE and BPaL (see Methods for dosing details). In all three regimens, we see substantial reductions of both granuloma and host CFU immediately starting treatment. In the highest-CFU granulomas and hosts, a precipitous drop in bacterial burden is followed by a rate of bacterial reduction at later time points in detailed granuloma-scale simulations^25,40^ (**Figure 4**A-F). We further compare our predictions to datasets from marmosets, a NHP model of active TB disease^40^, and indeed our predictions agree with those data for the most severe *HostSim* virtual infections, though all data are contained within the spread of our outcomes (see black dots in **Figure 4**). By contrast, the majority of our virtual cohort sterilize virtual infection within two months of HRZE and RMZE treatment, and within 1 month of BPaL treatment—comparable to new data from Mauritian cynomolgus macaques (green markers in **Figure 4**).

**Figure 4.**
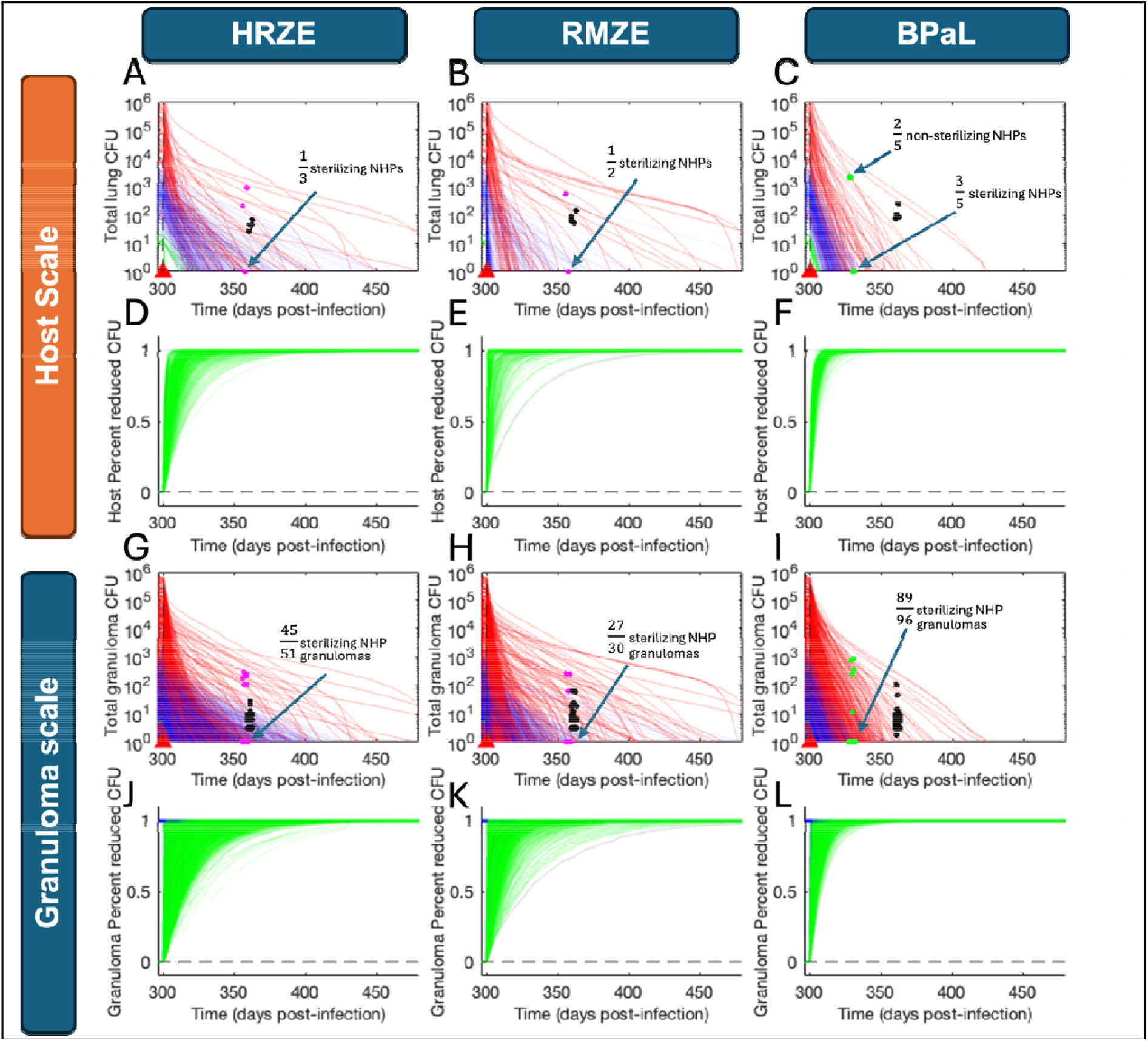
Efficacy of three simulated drug regimens, at the host and granuloma scale, compared with NHP data. We show simulated data from six months of treatment with HRZE (left column), RMZE (center column), or BPaL (right column). Treatment begins at day 300 post-infection (red triangles) to allow the infection dynamics in our virtual hosts to mature prior to intervention. We show raw CFU counts at both host-scale (A-C) and granuloma-scale (G-I). Both granuloma and host CFU curves are colored based on host infection classification prior to treatment—green for apparently-sterilizing, blue for subclinical and red for active disease. Black dots are CFU counts from marmosets, a NHP model of active disease^40^; magenta diamonds are 8-week total-lung (A,B) and lung granuloma (G,H) CFU values obtained from *Cynomolgus macaques* treated with HRZE and RMZE, respectively^25^; green dots are total lung (C) and lung granuloma (I) CFU measured from macaques treated with BPaL for 4 weeks prior to necropsy (Methods). For magenta and green markers, where applicable, the number of sterile (0 CFU) granulomas has been indicated to emphasize the difference between granuloma and whole-host sterilization frequencies and indicated overlapping markers for NHPs with similar CFU counts. We have added a random jitter to the x-values (maximum ±2 days) of each marker for visibility. Trajectories of Percent CFU Reduction impact scores (Methods section 4.9) are shown in time (D-F host-scale; J-L granuloma-scale), showing variation in both host and granuloma scale responses to treatment.

Next, we wanted to examine how much each virtual host was improved by antibiotic when compared to what they would have experienced without. To do this, we performed a MID-framework analysis (Methods) examining the percent reduction of Mtb within each host and granuloma. We find that for the vast majority of virtual hosts and granulomas, over 80% of CFU are killed within the first two months of treatment—regardless of regimen (**Figure 4**D-F host-scale, G-I granuloma-scale). This is consistent with the general consensus in the field: that the main challenge of TB treatment is sterilizing a smaller portion of persister Mtb.

### 2.5. HostSim results allow identification of mechanisms driving host response to treatment

We next identified the model mechanisms (key parameters) that most influence which hosts are most improved by treatment. We employ a two-step analysis to do this. First, we use a MID-framework analysis to generate quantified scores for improvement of infection outcomes on a per-host and per-granuloma basis (Methods Section 4.9, and **Figure 4**D-F and J-L). We measure four impact scores, collectively the *host-improvement scores*, that quantify efficacy of HRZE, BPaL and RMZE in killing Mtb within granulomas and hosts, specifically when compared to their CFU totals in the no-treatment scenarios^26^ (see Methods). Second, we use a multi-scale sensitivity analysis (Methods Section 4.10) to determine which features of the virtual host identity (i.e., host baseline parameters and specific granuloma parameters) drive variability in both total CFU in the control scenario (no treatment; CFU_control_) and each host improvement score. To do this, we perform the Partial Rank Correlation Coefficient (PRCC) global sensitivity analysis method (see Methods), which determines non-linear correlation values between parameter values (i.e., mechanism activity) and outcomes. Our PRCC results, summarized in (**Table 2**), show that variation in virtual treatment predictions is driven by several expected drivers that underpin drug efficacy.

**Table 2:**
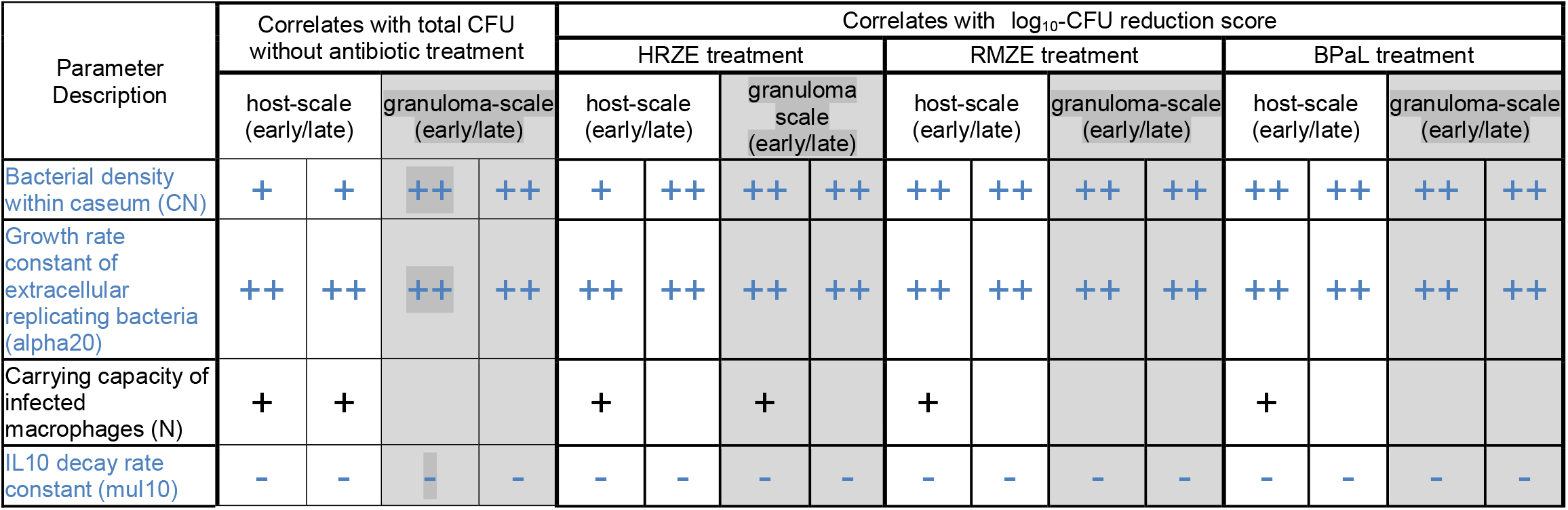
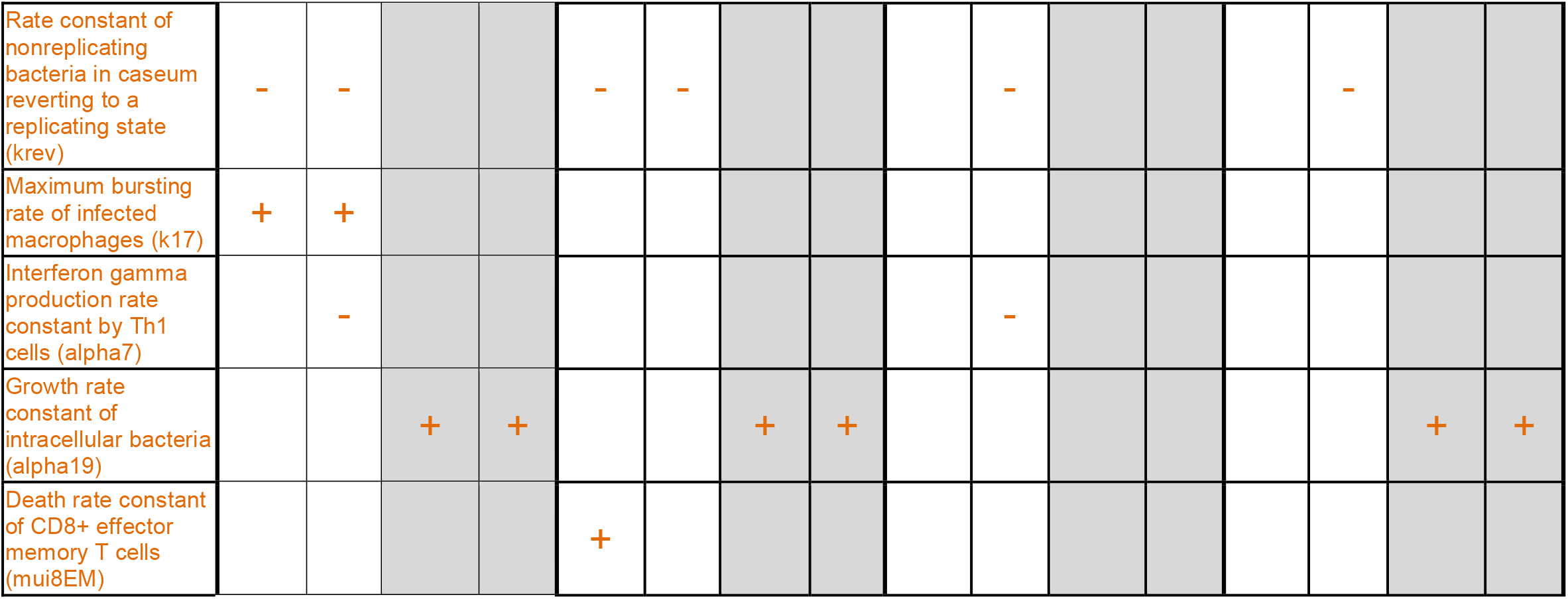
*HostSim* parameters that are significantly correlated to without-treatment CFU and log_10_-CFU reduction relative to the control during simulated HRZE, RMZE and BPaL treatment scenarios. Correlations are indicated if they are statistically significant at both the granuloma and host scales for at least 5 days (*p < 0*.*05*, Bonferroni corrected for multiple comparisons). Indicated columns refer to correlations deemed significant during the first 50 days of treatment (early) and after 50 days of treatment (late), respectively. The markers ++ and -- indicate that PRCC values lie between 0.4 and 0.6 or between -0.4 and -0.6, respectively. Similarly, + and - indicate that PRCC values lie between 0.2 and 0.4 or -0.2 and -0.4, respectively. Rows indicated in blue are common to all treatment scenarios as well as total control-scenario CFU. Orange rows indicate outcomes that are only significant when measured at one scale.

Of all four host-improvement scores, we found that log_10_ CFU reduction revealed the most CFU-influencing mechanisms that we group into three types of parameters: (i) those that influence treatment outcomes at the host level, and do so similarly for all three drug regimens, (ii) those that influence treatment outcomes at the host level, but differentially so between the three drug regimens, and (iii) those that primarily influence treatment outcomes at only one scale. We discuss each in turn.

Both CFU_control_ and log_10_ CFU reduction by all three regimens correlate with several expected granuloma and host baseline parameters (**Table 2** blue rows): IL-10 levels, rate of extracellular Mtb replication, and bacterial density within caseum. As expected, increased bacterial replication rates lead to larger Mtb populations.

Similarly, in virtual granulomas that have larger fractions of Mtb that survive macrophage death and become trapped within caseum, there are larger total Mtb-counts. Finally, IL-10 has been observed to impede adaptive immunity to pulmonary Mtb infection, resulting in higher CFU levels^53^, although the dynamics of IL-10 involvement are more complex^18,20,54^. All three of these mechanisms result in a larger Mtb population to be killed by antibiotics, proportionally increasing log_10_ CFU reduction scores. However, mechanisms characterized by these parameters do not differentially affect the efficacy of our three simulated antibiotic regimens.

The collection of parameters driving the total CFU without treatment account for most of the parameters that correlate to the three other host-improvement scores: non-replicating CFU reduction (**Supplemental Table 1**), percent reduction of total CFU (**Supplemental Table 2**), and percent reduction of non-replicating CFU (**Supplemental Table 3**). That is, most parameters that drive host-improvement score with any regimen also drive the number of CFU without treatment. This is because CFU-reductions are larger in granulomas with higher initial CFU levels, whereas hosts with low-CFU burdens sterilize faster—leading to higher percent-reduction scores. (The two exceptions, where parameters significantly correlate to host-improvement scores but not to total CFU count, are discussed below as parameters that differentially influence regimen efficacy.) We observe that EBA-like scores (log_10_ CFU reduction and log_10_ non-replicating CFU reduction, see Methods) are positively correlated with parameters that are themselves correlated to total bacterial burden during the course of the treatment. This contrasts with scores based on percent-reduction of total and non-replicating Mtb counts within hosts and granulomas (see Methods), which are negatively correlated to the same parameters only early in treatment.

**Table 3:** Hill curve parameters for each drug and each Mtb location for PD model.

| Drug | Mtb location | $E_{\max}$ (1/day) | $C_{50}$ (mg/L) | $h$ | Reference |
| --- | --- | --- | --- | --- | --- |
| Isoniazid | Intracellular | 0.66 | 0.092 | 1.05 | 86 |
|  | Extracellular | 1.18 | 0.012 | 1.8 | 86 |
|  | Nonreplicating | 0.3 | 1.86 | 1.29 | 87 |
| Rifampicin | Intracellular | 1.73 | 0.6 | 0.4 | 88 |
|  | Extracellular | 6.48 | 0.081 | 2.14 | 88 |
|  | Nonreplicating | 0.69 | 4.14 | 1.55 | 87 |
| Pyrazinamide | Intracellular | 0 | 0 | 0 | 89 |
|  | Extracellular | 0.27 | 13.66 | 0.94 | 89 |
|  | Nonreplicating | 61.92 | 1124.73 | 1.04 | 87 |
| Ethambutol | Intracellular | 1.24 | 0.5 | 2.73 | 90 |
|  | Extracellular | 1.31 | 0.5 | 3.45 | 90 |
|  | Nonreplicating | 0 | 0 | 0 | 91 |
| Moxifloxacin | Intracellular | 0.53 | 6.9 | 5.39 | 92 |
|  | Extracellular | 1.07 | 0.06 | 4.99 | 92 |
|  | Nonreplicating | 48.96 | 0.83 | 4.52 | 87 |
| Bedaquiline | Intracellular | 0.92 | 0.12 | 4.41 | 93 |
|  | Extracellular | 61.92 | 2.54 | 1.67 | 93 |
|  | Nonreplicating | 0.56 | 5.14 | 0.74 | 87 |
| Pretomanid | Intracellular | 0.46 | 0.07 | 2.62 | 94 |
|  | Extracellular | 0.94 | 0.21 | 0.97 | 95 |
|  | Nonreplicating | 0.56 | 5.06 | 1.33 | 96 |
| Linezolid | Intracellular | 0.43 | 0.3 | 2.6 | 94 |
|  | Extracellular | 0.76 | 0.51 | 2.08 | 97 |
|  | Nonreplicating | 0.34 | 1.35 | 0.94 | 87 |

By contrast, some mechanisms differentially influence regimen efficacy. Reduced host baseline IFNγ production rates correlate with improved RMZE log_10_-CFU reduction of total CFU Mtb, suggesting that RMZE could be a better regimen for those patients who have lower levels of IFNγ. As MXF and RIF have relatively poor penetration into caseum^55^, this may suggest that higher levels of IFNγ result in lower-CFU yet more caseated granulomas on average. Moreover, during the first 50 days of treatment, granuloma-scale log_10_-CFU-reduction scores in HRZE-treated hosts correlate with higher carrying capacity of Mtb within infected macrophages. In that same scenario, hosts with larger baseline intracellular Mtb growth rates correlate with improved HRZE and BPaL CFU-reduction scores. Next, log_10_ reduction of CFU is (but not CFU_control_) increased during HRZE treatment with higher death rates of CD8^+^ Mtb-specific effector memory T cells, suggesting that CD8^+^ Mtb-specific T-cell differentiation to effector states may play a significant role in understanding heterogeneity in host response to treatment. Finally, increased rates of TNF-driven recruitment of macrophages to granulomas negatively correlated to the log_10_ reduction of non-replicating CFU (but not control-scenario non-replicating CFU) during BPaL treatment. Notably, if one could potentially characterize activity level of these mechanisms within individual patients, these results could provide insights on personalizing regimen choices between HRZE, RMZE, or BPaL.

Finally, some mechanisms driving CFU_control_ and/or CFU-reduction scores have parameter values that are sensitive only at one scale (**Table 2** orange rows). At only the host scale, we see negative correlations between both CFU_control_ and/or CFU-reduction scores and the rate of non-replicating bacteria removed from caseum; this mechanism is included in our model to capture neutrophil-driven movement of Mtb phagocytized within caseum and carried to the granuloma cellular area^16,56-58^. Also at the host scale, we see single-scale correlations: the maximum bursting rate of infected macrophages, CD4^+^ T-cell production of IFN-γ, and presence of CD8^+^ T cells; all of which are both known to be influential processes in determining host outcomes^4^. The fact that the host-baseline parameters correlate with host outcomes, but the granuloma-specific parameters do not, indicates that the impact of these mechanisms on total lung CFU may be obscured by the noise of within-host heterogeneity, while the host’s baseline activity of these mechanisms still has a measurable impact on whole-host outcomes. Conversely, growth rate of Mtb within macrophages has a significant impact on granuloma CFU, although the host baseline parameter value is insufficient to predict the host-scale CFU. Together, these results indicate that the impact of some mechanisms are principally driven by the normative activity of a mechanism within an individual, while others are driven by intra-host granuloma heterogeneity.

### 2.6 *HostSim* drug rankings correlate well to multiple previous regimen-ranking studies

We have demonstrated that our *HostSim* antibiotic treatment model captures the complex multi-scale process of antibiotic TB treatment, recapitulates a very broad range of responses at the host-scale, and captures mechanisms underpinning a small number of well-studied regimens. We next want to ensure that we capture population-scale trends including the relative efficacy of multiple drug regimens. To this end, we further validate and predict with *HostSim* by generating antibiotic efficacy rankings of different sets of antibiotic regimens similar to prior studies^40,50,59^. Our ranking methods include 1) virtual 8-week measurement of apparent-sterilization and 2) measurement of area under a regimen’s sterilization curve (AUSC; see Methods for both), and we examine those rankings at the granuloma and whole-host scales.

For all regimen sets (i-iii) (see **Table 6**, Methods), we are able to reproduce regimen efficacy rankings from *HostSim* that agree well with those from literature (i.e., significant Spearman correlation with *R* >0.75) ^40,50,59^. Using a straightforward measurement of 8-week apparently-sterilizing virtual hosts, we produce similar results to clinical rankings published for regimen set (i), a collection of various HRZEM-containing regimens^50,59^ (**Figure 5**A). Regimen set (ii), a collection of BDQ- and MXF-containing regimens, has been previously studied in marmosets (animal model of progressive TB disease^60^). Thus, in order to directly compare *HostSim* to rankings derived from marmoset data, we generated rankings based only on high-CFU virtual hosts and granulomas (see Methods). By comparing the same AUSC-based ranking approach to marmoset rankings as in our previous work^40^, we find that *HostSim* rankings of high-CFU granulomas are similar to the CFU-based rankings obtained from CFU reduction in marmoset granulomas (**Figure 5**B). Finally, our granuloma-scale 6-month AUSC-based rankings agree with rankings given by our detailed granuloma-scale model, *GranSim*^*25*^, which were computed using the same method (**Figure 5**C). Together, these results suggest that *HostSim* captures relative efficacy of antibiotic regimens for TB treatment in the context of drug-susceptible Mtb infection.

**Figure 5:**
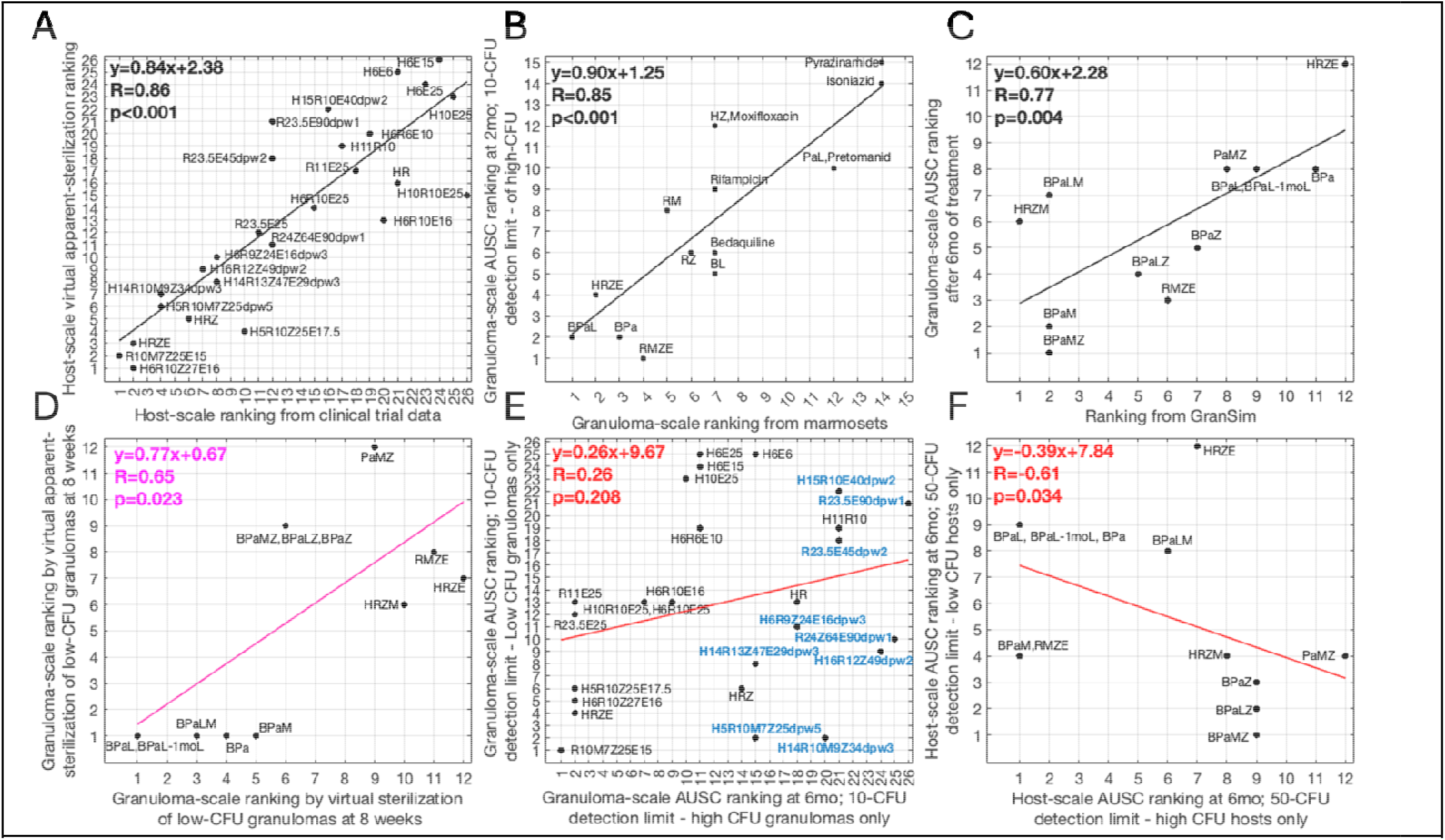
Comparing ranking methodologies of drug regimens at both host and granuloma scales. In each panel, we show a Spearman rank correlation between two ranking methods for a set of antibiotic regimens. We have colored the best-fit line equation, spearman’s rho, and p-value for each panel based on the fit—magenta if R<0.7, red if R<0.3. Our indications of regimen set (i-iii) are in reference to the regimen sets from Methods **Table 6**. Note that we have normalized each ranki g, particularly those from prior work, so that in all panels better-performing regimens have lower ranks. (A-C) We show validation of our drug model by reproducing previously-established rankings, and (D-F) qualitative observations when comparing differ nt ranking methodologies within *HostSim*. (A) Ranking of regimen set (iii) based on 8-week host-scale apparent-sterilization agree with clinically-established rankings^50,59^. (B) Rankings of regimen set (ii) based on high-CFU *HostSim* hosts using 2-month AUSC ranking method agree with Marmoset rankings established in previous work^40^. (C) Rankings of regimen set (i) based on granuloma 6-month AUSC-rank agree with identically-computed ranking method from *GranSim*^40^. (D) Poorly-correlated rankings of regimen set (i) are computed by considering 8-week apparent-sterilization versus sterilization. (E) We show apparent-sterilization 6-month AUSC-based rankings of drugs from regimen set (iii) when the cohort is restricted to only high-CFU (y-axis) or low-CFU (x-axis) virtual granulomas. Drugs with less frequent dosing (indicated in blue) have substantially worse ranks for high-CFU granulomas. (F) In regimen set (i) apparent-sterilization AUSC ranks at 6-months do not correlate between high-CFU and low-CFU granuloma at host scale.

### 2.7 Cohort bacterial burden and detection limit influence drug rankings

Finally, we wondered whether the choice of ranking method and accuracy of associated measurements affects various antibiotic regimen rankings, and whether our validated treatment model has sufficient detail to address this. Each ranking method tested here has been developed as a means of characterizing sterilizing potential for each regimen; differences in each method represent different experimental or clinical trial study designs. Broadly, we calculate two main types of rankings: apparent-sterilization rankings (an estimation of simulated solid culture conversion) and area under the sterilization curve (AUSC)-based rankings (see Methods). We generated these results by differently-calculating rankings for regimens that treated the same virtual cohort examined in the previous sections, giving a single “ground-truth” of the infected population underpinning these rankings. We perform these rankings at the host-scale (i.e., using total-lung CFU data) and granuloma-scale rankings (i.e., pooling data from each virtual primary granuloma). To examine the impact of detection threshold, we draw a distinction between analyzing simulations with respect to virtual host *sterilization* (CFU < 0.5 within the model) and virtual host *apparent-sterilization* (CFU < 10 in a granuloma, or CFU < 50 in a whole-host; see Methods). Finally, to examine the impact of inclusion criteria, in some analyses we subset our virtual cohort into high-CFU and low-CFU patients and granulomas (see Methods).

For regimen set (i) (see **Table 6**), we find relatively poor correlation () between rankings of treatment efficacy generated by measuring fractions of simulated low-CFU granuloma sterilization vs. apparent-sterilization at 8 weeks (**Figure 5**D). By contrast, we find a high correlation between rankings generated by measuring the same fractions in sets (ii) and (iii) (both with *R* > 0.95, *p* <0.001, see **Supplemental Figure 1**). The poor correlation in regimen set (i) suggests that some regimens may be more likely to permit undetected Mtb persistence within hosts and granulomas—enough to substantively affect drug rankings. This finding is corroborated in literature: according to the sterilization-based rankings of regimen set (i), HRZE is less efficacious than RMZE or HRZM, which is consistent with mouse^42,49^ and macaque studies^25^. However, according to apparent-sterilization rankings, HRZE performed better than RMZE but worse than HRZM (**Figure 5**D).

There are multiple aspects of experimental design that affect our simulated rankings, including (1) heterogeneity of the mechanism of action in each regimen set, (2) dose frequency, and (3) inclusion criteria for the virtual cohort. We see a lack of clear correlation between virtual apparent-sterilization-based rankings computed from high-versus low-CFU granulomas (**Figure 5**E). The lack of correlation in regimen set (iii) may be caused by the diversity of treatment frequencies within the regimen set, as regimens with less frequently administered dosages tend to have better rankings for low-CFU granulomas than high-CFU granulomas (**Figure 5**E blue regimens). Moreover, regimen set (i) contains a diverse set of antibiotics in terms of their mechanisms of action (e.g., ability to penetrate caseum and substantially different PD parameters by spatial region), hence their rankings depend highly on bacterial burden. This results in a lack of correlation between the rankings of high- and low-CFU granulomas (**Figure 5**F). Furthermore, our rankings based on low- and high-CFU granulomas agree with the stratified approach that claimed MXF-containing regimens may replace HRZE for patients with minimal disease^61^, as MXF-containing regimens are ranked higher in low-CFU granulomas (**Figure 5**E and F). Together, we see that a variety of experimental design choices in a description of sterilization-capability can substantially affect conclusions drawn from regimen ranking analyses.

## 3. Discussion

The question of how to improve TB treatment 100 years after antibiotics were discovered and over 50 years since TB has been treated with a 4-drug regimen still remains unanswered. Human trials are costly, and with the discovery of new drugs even animal models are insufficient to compare the many regimen possibilities. Simulations offer a novel approach to help narrow the design space of possible regimens. To this end, we present results from a multi-scale model of pulmonary Mtb and multi-drug antibiotic treatment regimens. *HostSim* recapitulates multiple features of host-pathogen interactions within heterogeneous Mtb-infected lesions in the lung across physiological scales ranging from cellular/molecular to tissue to organ and to whole host. We can generate a virtual cohort and thus have the unique ability to test the same set of virtual hosts with multiple antibiotic regimens for direct apples-to-apples comparison—not possible in any other system. We demonstrate here that *HostSim* captures the spectrum of TB outcomes, matches clinical EBA and experimental NHP CFU measurements, and can produce mechanistically-sound virtual clinical trials that reproduce existing regimen rankings.

In the absence of treatment, our model predicts that only one of our 500 virtual Mtb-infected hosts (either with subclinical infection or active disease) completely clears infection (sterilization). In some virtual hosts, CFU levels reach below-detectable levels; these hosts often sequester surviving Mtb within caseum. Poorly-controlling (high-CFU) granulomas in our model are associated with large intra-macrophage Mtb populations. Our model predicts that parameters controlling bacterial growth, Mtb density within caseum, levels of IFN γ and IL-10, and infected macrophage-Mtb interactions are predictive of overall bacterial burden. These findings suggest a dual role of IL-10, as it has been observed to correlate with faster sterilization rates^18,20,54^; this suggests that inflammation may worsen CFU burden in some granulomas while remaining important for complete sterilization. In virtual hosts with active TB, high-CFU granulomas have macrophages that, on average, permit high levels of intracellular bacteria and lack support from sufficient CD4^+^ or CD8^+^ T cells to either activate macrophages or induce apoptosis of infected macrophages, thereby providing a larger replicative niche for Mtb. By contrast, low-CFU simulated granulomas have sizeable non-replicating Mtb populations within caseum and maintain low levels of intracellular bacteria (blue trajectories in **Figure 1**), which suggests that confining Mtb in caseum is representative of well-controlling granulomas.

Our *HostSim* model includes a detailed PK/PD model reporting individual drug concentrations throughout host blood, lung tissue, and within each granuloma. We represent regimens that include both front-line and last-resort treatments for TB, specifically HRZE (aka RIPE, consisting of INH, RIF, PZA, and EMB) as well as multiple antibiotics used to treat drug-resistant infection (BDQ, PTM, LZD, and MXF). Most humans treated for drug-susceptible TB sterilize by 4 months, but a smaller percentage take up to 6 months^41,62^, similar to what we observe in our virtual cohort (**Figure 3**). Our model also includes the influence of drug-drug interactions. In simulations of treatment with well-known regimens (e.g., HRZE, RMZE and BPaL), we observe biphasic CFU reduction in both granulomas and hosts, where CFU levels drop quickly early during treatment and more slowly later in treatment (**Figure 4**), which has been observed in prior Mtb studies^21,63^. We observe this in our simulations when a sparse number of surviving Mtb are distributed among a large number of infected macrophages, leaving a rich replicative niche. During treatment, biphasic reduction also has been observed in HIV viral load^64^, another pathogen with an intracellular replicative niche.

Importantly, once we simulate antibiotic regimens, some method of ranking outcomes is needed to assess relative efficacy. We developed predictive regimen rankings based on performance over each spatial scale. We based these ranks on two measures of drug sterilizing potential: proportion of hosts that are apparently-sterilizing by eight weeks post-treatment, and area under the sterilization curve (see Methods). We find that by several metrics assessing overall CFU reduction, that BPaL and similar BDQ-containing regimens are among the top performers when compared against multiple drug regimens (**Figure 5**). However, there is not a clear winner among drug regimens. RMZE performs well in short-course trials when only compared against other HRZEM-containing regimens with varied dose frequencies, whereas RMZE struggles in comparison to BDQ+PTM-containing regimens when only examining high-CFU hosts and granulomas. By sterilization time (AUSC ranking method), we predict that BPaMZ is the best out of 12 regimens (set (ii), see **Table 6**) when applied to low-CFU hosts, yet performs poorly for high-CFU hosts (**Figure 5**). To summarize, we find that regimen efficacy rankings depend on the ranking methodology.

Overall, we find that a single definition or metric for measuring drug efficacy, even when characterizing an intuitive feature (e.g., “CFU sterilizing potential”) is insufficient to appropriately characterize available treatment options for TB. We find that when antibiotics have very different actions, even the distinction between sterilization and apparent-sterilization can render poorly-correlated conclusions. Our results highlight the potential non-interchangeability of superficially-similar ranking methods—e.g., the importance of clear and consistent descriptions of inclusion criteria and experimental limitation during clinical trial design, and the importance of multi-ranking consensus or context specificity when endorsing one treatment regimen over another.

The discrepancies we identify between ranking methods and overall conclusions may explain conflicting results seen between finings in the literature. For example, in the ReMOXTB study, MXF-containing regimens demonstrated a better sterilizing power but lacked long term efficacy due to relapse rates resulting from undetected Mtb^41^. Mouse studies provided evidence that HRZE is significantly less potent than HRZM and RMZE^42,49^ as shown in our rankings based on host sterilization (**Figure 5**D). However, clinical trials have inconclusive results for the performance of HRZE compared to RMZE and HRZM^41,65,66^. This agrees with our rankings based on culture negativity versus sterilization, where HRZE, RMZE and HRZM have very close rankings and HRZE is not significantly worse than RMZE. Based on our sensitivity analysis, we predict that hosts that are more prone to inflammation will have lower-CFU yet more caseated granulomas. This will affect the relative efficacy of RMZE as this regimen has lower levels of caseum penetration^55^. This merits an in-depth exploration on the role of inflammation on caseum accumulation. Moreover, because virtual host baseline levels of IFNγ correlate with reduced log_10_ CFU reduction score of RMZE, it may be that host-level IFN γ concentrations—perhaps measurable from blood during an IGRA^67^ or a similar ELISA^68,69^ test—may be a predictor of RMZE efficacy.

Unintended measurement bias may also extend to the assessment of individual host or granuloma response to an antibiotic regimen if a single “key outcome” is taken as an index for regimen efficacy. To explore which host mechanisms correlated with improved host-response to treatment via host-improvement scores defined *a priori*, we used a MID framework analysis, in which virtual hosts serve as their own controls^26^. We find several parameters that correlate with improved host-response to all of three regimens: higher Mtb density with caseum, faster growth rate of extracellular Mtb, and longer persistence of IL-10—all intuitively related to more severe TB disease. Moreover, we find several parameters that improve host-response scores of some regimens but not others. For example, we predict that higher growth rates of Mtb within macrophages statistically significantly improves the host response to HRZE and BPaL, but not to RMZE (see **Table 2**). These results may seem counter-intuitive, but it becomes clear when scrutinizing the definition of improvement. Hosts that have the highest CFU burdens harbor more Mtb to kill, especially in regions outside of caseum. Virtual hosts with more severe TB have more Mtb that is not confined to caseum and in larger quantities, resulting in more rapid CFU reduction in non-caseated niches and, consequently, a more-impressive CFU-reduction score. Conversely, patients that control infection well and sequester Mtb to caseum will, comparatively, have lower CFU reduction regardless of whether their non-replicating CFU are killed or not. This highlights how CFU reduction alone does not well-characterize the ability of antibiotic regimens to eliminate populations of persister bacteria.

While *HostSim* does include antibiotic treatment, drug-drug interaction, T-cell priming, granuloma dissemination, and caseum necrosis, additional biological details may further improve our simulation platform and are the topic of future work. First, we do not yet include explicit representation of lymph node infection, which is known to correlate with more severe disease states^70^. Incorporation of bacterial resistance will be important to understanding the effects of poorly adhering to antibiotic drug regimens^34^. Moreover, we assume that our bacterial strains are entirely susceptible to each of our drugs and did not take drug resistance into account. In addition to human granulomas, we have calibrated our model to match datasets from non-human primate and rabbit granulomas. While both of these animal models have similar features to human TB, more data from lymph node infection states during Mtb infection would provide more nuanced predictions. Lastly, post-treatment relapse is another significant aspect to assess treatment success that will require modeling factors that influence long-term T-cell activity^71^. We plan to refine our models and include these crucial components for more in-depth virtual preclinical trials.

## 4. Methods

In the subsections below, we detail *HostSim*, our multi-scale mechanistic model of TB progression and treatment. We also provide a detailed description of experimental methods used for the measurement of BPaL efficacy in Mauritian cynomolgus macaques.

### 4.1 Overview of *HostSim*, our multi-scale representation of Mtb infection and drug treatment

In this study, we use the next-generation version of our computational model *HostSim*^*26*,*27*,*72-75*^ to analyze virtual cohort clinical trials across multiple scales (**Figure 6**). *HostSim* allows us to simulate whole-host outcomes of pulmonary Mtb infection within a single virtual host with multiple lung granulomas (**Section 4.2**). We assume that a single collection of parameter values defines the *virtual identity* of any virtual subject (host, granuloma, etc.), and we generate a stratified, diverse virtual cohort by using a Latin hypercube sampling method (**Section 4.3**) using parameter ranges calibrated via our calibration protocol, CaliPro (**Section 4.4**). We create a *virtual cohort* of 500 virtual hosts, each containing 13 primary granulomas (for a total of 6500 primary granulomas). This follows since data from humans and NHPs suggest that primates on average have approximately 13 granulomas at the time of infection with the possibility of more disseminating during infection. This assumption reflects a specific low-dose inoculum; our model can be updated to accommodate more or less numbers of granulomas.

**Figure 6:**
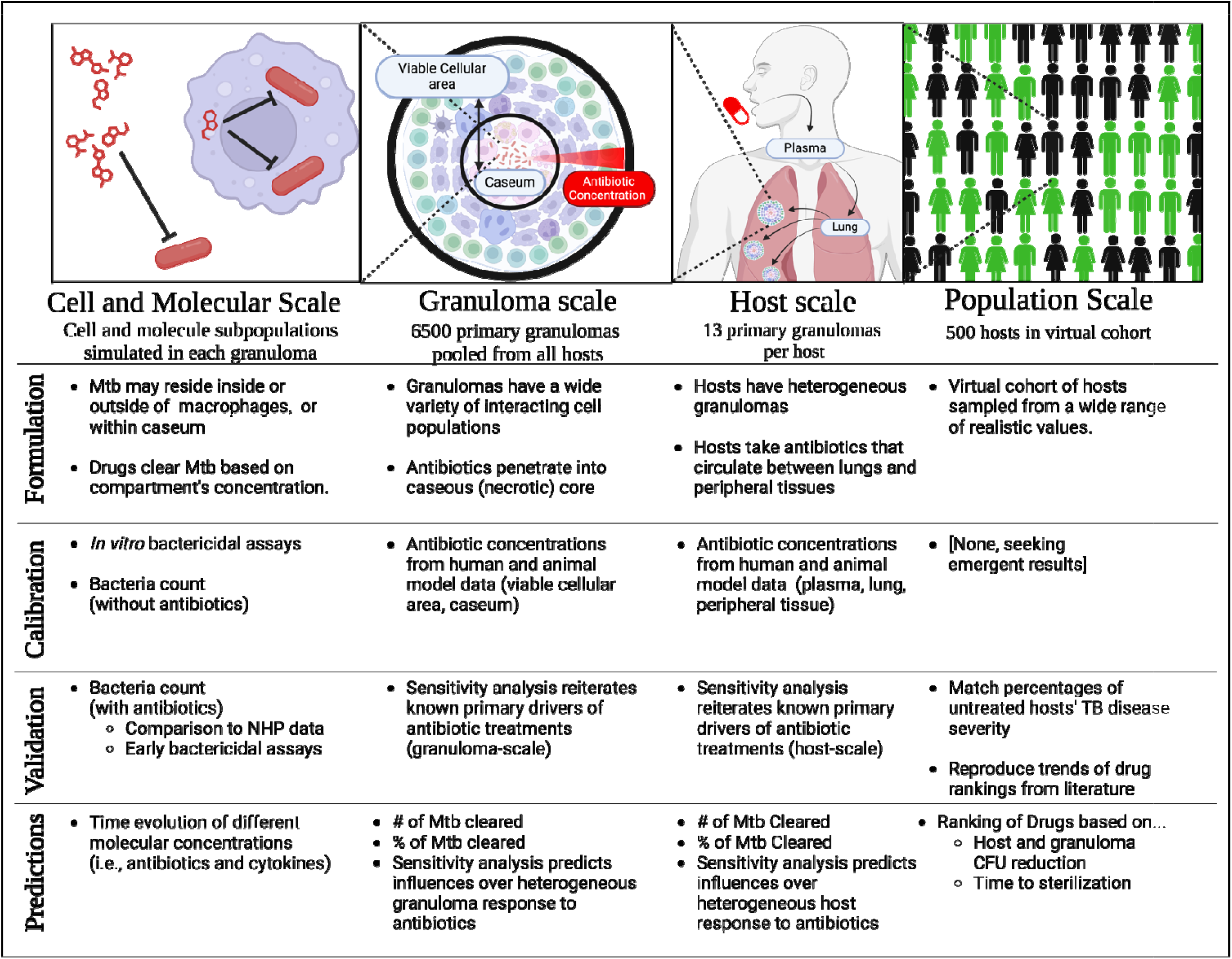
*HostSim* captures multiple spatial and temporal scales. Top-to-bottom and by-sc*a*le, we summarize key features of our model, specify model calibration metrics, identify validation targets, and outline the measurements that our model will predict. (Figure created with bioRender.com.)

In this work, we extend *HostSim* by incorporating PK/PD and drug-drug interactions to study the dynamics of eight well-known and experimentally studied anti-Mtb antibiotics (**Section 4.5**) within multi-drug regimen (**Section 4.6**).

To analyze this model, we use and develop several unique multi-scale approaches. We define our virtual measurements of drug efficacy (**Section 4.7**) and also create multiple drug ranking algorithms for comparison (**Section 4.8**). We discuss other mathematical tools that we use to further investigate our virtual cohort, including our multi-scale interventional design (MID) framework^26^ (**Section 4.9**) and sensitivity analyses^76^ (**Section 4.10**).

### 4.2 Immune-Mtb interactions in *HostSim*

The response of lung granulomas to antibiotic treatment is heterogeneous both between hosts and within an individual host^4^. Consequently, we need a whole-host computational model, including simultaneous treatment of multiple granulomas, to further investigate the impact of drug regimens. To this end, we create a next-generation version of *HostSim*, our previously-published whole-host model of bacterial-immune interaction of virtual hosts infected with Mtb^26,27^.

Briefly, virtual hosts in *HostSim* have a lung compartment with multiple granulomas, a draining lymh node compartment, and a blood compartment. Each granuloma in the lung compartment can be considered an ‘agent’ in the computational model. Granulomas are comprised of different cellular subpopulations and proteins, including immune cells, Mtb, and cytokines, and are represented as a system of ordinary differential equations (ODEs). Immune cell types include macrophages and CD3^+^ T-cells (both CD4^+^ and CD8^+^) in different stages of differentiation. We subdivide Mtb populations within each granuloma into intracellular within macrophages (), extracellular within granuloma spaces (), and caseum-trapped non-replicating () subpopulations. Immune cells within the model respond to bacteria by releasing inflammatory cytokines such as tumor necrosis factor alpha (TNF) and Interferon gamma (IFN). Mtb-infected granulomas send antigen-presenting cells (APCs) to lung draining lymph nodes (LDLNs) that prime and clonally expand Mtb-specific effector CD4^+^ and CD8^+^ T cells. Once activated, effector T cells migrate into blood where they can be recruited into granulomas in response to recruitment signals. See **Appendix A** for a complete list of *HostSim* equations.

We update how we track bacterial dynamics within *HostSim*, and include the addition of free Mtb antigen, not just whole Mtb. We include this in the representation of lung-LN interaction by tracking antigen within the LN compartment. Previously, our model of antigen presentation tracked infected macrophages as a proxy for amounts of bacterial antigen; we now explicitly represent a calculated quantity of Mtb antigen within granulomas traffics to LDLNs within dendritic cells and macrophages. Level of antigen is tracked by accumulation when Mtb dies and antigen degrades slowly over time. For details on how antigen affects T-cell priming, see **Appendix A**.

### 4.3 - Parameter Selection, virtual cohort construction, and virtual host identity

We generate virtual cohorts in *HostSim* using parameter sets that capture both inter- and intra-host heterogeneity. Non-human primates exposed to small infecting inocula (<40 CFU) have been observed to form up to two dozen granulomas when observed for 4 weeks^77,78^. To simplify our model, and with the understanding that some virtual granulomas never develop above a 50CFU detectability threshold, we seed each of our virtual hosts in our virtual cohort with 13 primary granulomas (representing a low-dose inoculation for a total of primary granulomas when pooled across-hosts). During the course of virtual infection, granulomas may disseminate and form a granulomas within each host, becoming more likely at higher CFU levels (see **Appendix A**). For each parameter value, we select a *host baseline parameter value* via the Latin hypercube sampling (LHS) method (**Figure 7**A, see **Appendix A** for a complete listing of parameters and their ranges). To generate values to parameterize ODEs for each granuloma within a host (e.g. pulmonary granuloma parameters and granuloma-scale PK parameters), we sample from a host-baseline-informed normal distribution. Specifically, we use the baseline host value for each granuloma granuloma parameter as the mean for a Gaussian distribution that we further truncate by the parameter range. We let the standard deviation for each Gaussian be to promote intra-host heterogeneity and best match the distribution of sterilizing granulomas per host. We use log-normal distributions similarly to sample parameters whose ranges span more than two orders of magnitude.

**Figure 7:**
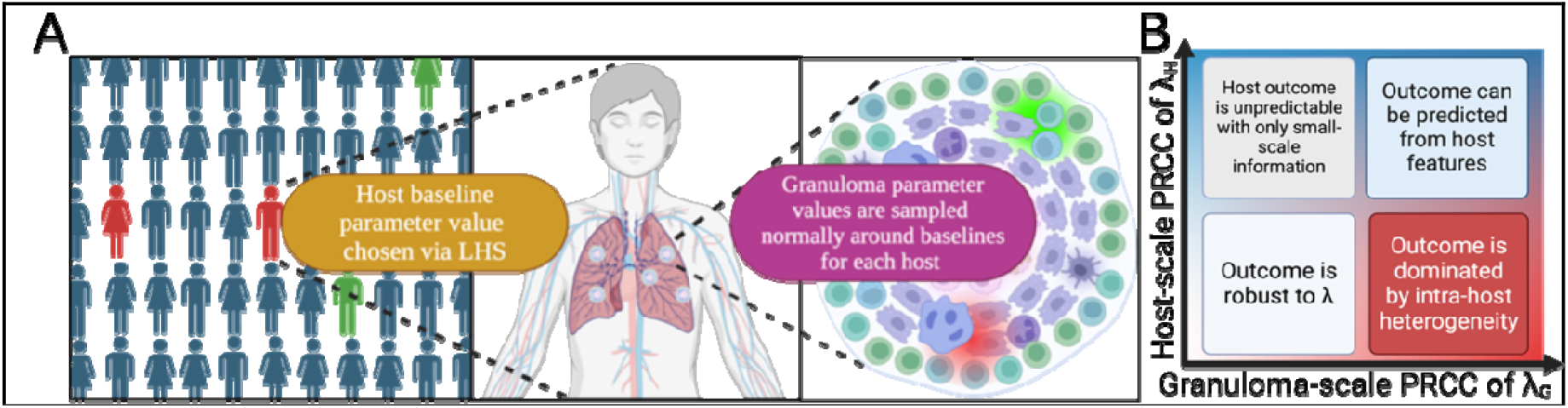
Intra-host heterogeneity and multi-scale sensitivity analysis. (A) We generate virtual granulomas in *HostSim* by sampling baseline parameter values for each host from calibrated ranges. Individual granuloma parameter values are sampled normally around baselines per-host to capture intra-host heterogeneity. Granuloma- (host-) scale outcomes can be analyzed via sensitivity analyses as functions of individual parameter (baseline) values. (B) *HostSim* produces output at two spatial scales, host [H] and granuloma [G]. Sensitivity analysis can determine the influence of any parameter (and thereby mechanism, e.g., λ) over model outcomes via a partial rank correlation coefficient (PRCC) (see Section 4.10). By comparing sensitivity analysis results that capture information about biological processes at each scale (e.g. bacterial burden capturing infection severity), we determine whether host baseline parameter values (, blue) can predict granuloma-scale outcomes, or if granuloma-scale information (, red) is required.

We refer to the collection of host baseline parameters, granuloma-specific parameters, and initial conditions of all host features (e.g., initial immune cell counts) as a *virtual host identity*, or a virtual host. It is important to note that this is distinct from dosages and timings of specific antibiotics. The virtual host identity includes everything that is pertinent to characterize a single individual with respect to mechanisms represented by *HostSim* that can be transferred between different scenarios. By sampling multiple virtual host identities, we create a heterogenous population within as our virtual cohort. When we refer to the same virtual host being treated HRZE versus BPaL, we mean that we have two simulations that have identical host baseline values, and identical granuloma-scale parameters. and only differ with respect to parameters unique to the intervention (i.e., drug regimen or other interventions). Note that our virtual hosts are similar to digital twins: a virtual host is a digital twin when its parameter ranges are calibrated to capture the outcomes a specific real-world counterpart^79,80^. (In events that unique digital twins are not identifiable, then a collection of virtual hosts most like a specific patient can be used as a set of *digital partners*^*26*^.)

### 4.4 Model calibration using CaliPro

To ensure that our model captures known primate TB dynamics, we perform a rigorous calibration to match *HostSim* outputs to a variety of different datasets and data types. In our previous work^26,27^, we calibrated *HostSim* to published datasets of NHP data^30-33^. To perform calibration we used *CaliPro*, a calibration tool that we developed to narrow parameter ranges while preserving biological features of a model^81,82^. Briefly, using *CaliPro* we take a wide range of parameter values as an input, evaluate the model using many parameter values sampled from that range, and then reject “failed” parameter values to narrow the range and better capture the “passing set” of granulomas. *CaliPro* is flexible in that the user may define a simulation run as “failed” based on biological heuristics that have been measured in the system.

Similar to previous work^26^, we define our pass set based on two types of datasets that we calibrate *HostSim* to–experimental and synthetic. Our experimental datasets include total Mtb CFU per granuloma, T-cells per granuloma, and T-cell-to-macrophage-count ratios, all taken from existing NHP datasets^30-33^. The synthetic datasets include granuloma caseum volume, macrophage subpopulation ratios, and Mtb subpopulation ratios generated from *GranSim*^16-19,21-23^. We consider a simulation as passing if its trajectories stay within one order of magnitude of all datasets simultaneously. We simulate 500 virtual hosts to generate a diverse virtual cohort and to capture a full spectrum of TB outcomes. Details of our use of *CaliPro* for this model are in **Error! Reference source not found**..

### 4.5 Representation of antibiotics within *HostSim*

We capture both relevant PK and PD for antibiotics detailed in the following subsections. Briefly, we predict concentrations of each drug in each spatial region of a virtual host. This PK is complicated by the kinetics of drug penetration into caseum within the dense granuloma core, which results in an inhomogeneous spatial distribution of antibiotics *within* caseum^21^; we implement a submodel to account for this distribution. Representation of PD in the model allows for Mtb to be killed at concentration-dependent rates that we capture by previously-calibrated dose-response curves. For all drugs that exceed a minimal inhibitory concentration, we assign kill-rate constants based on *in vitro* data, that are further modified by transcriptomically-informed predictions of drug-drug interactions^24,83^. Finally, we describe a mechanism that coarse-grains the distribution of how infected macrophages are cleared of Mtb as the intracellular bacteria population is reduced. We present the details of these steps below.

#### 4.5.1 Pharmacokinetics (PK) modeling

Using this next-generation version of *HostSim*, we give virtual hosts a simulated drug regimen—consistent doses of various drugs administered at prescribed intervals. Drugs are taken up into blood over time, then from blood they diffuse into both lung and peripheral tissues. From lung tissue, drugs permeate into the viable cellular regions of each granuloma and, finally, some drugs (but not all) are able to penetrate into caseum. We represent this process in *HostSim* by the inclusion of a new host-scale, blood-lung PK compartment given by the following system of equations:

Transit compartment 1,

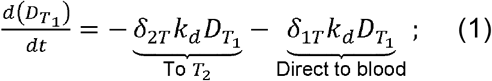

transit compartment 2,

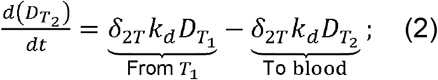

drug concentration in peripheral tissue,

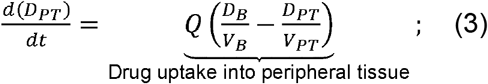

drug concentration in blood,

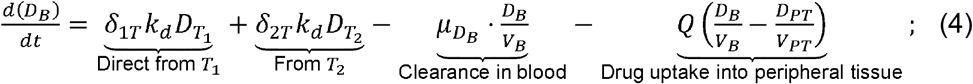

and total drug concentration in the lung,

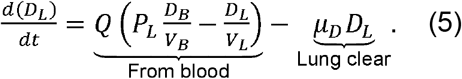

We pulse the prescribed dose into transit compartment 1 (with concentration 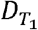) every 24 hours. We use a second transit compartment, with drug concentration 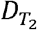, to capture the PK behavior of INH, PTM, RIF; whereas BDQ, EMB, LZD, MXF, and PZA only require one transit compartment^22,40^. We manage this by setting one of δ_2*T*,_ δ_1*T*_ to 1 and the other to 0 as appropriate. After passing through the transit compartments, the drug passes into the blood compartment (concentration *D*_*B*_) where it permeates into lung tissue (concentration *D*_*L*_) or be exchanged with peripheral tissue (concentration *D*_*PT*_) We transition between mass concentration (mg/kg) and volumetric concentration (mg/L) using tissue volume distributions (*V*_*PT*_). The parameters *k*_*i*_ are rate constants determining how quickly drug moves through the transit compartments. We can use the lung concentration (5) of each drug to predict the amount of drug within the virtual host’s granulomas via the following equations:

Drug amount (in mg) in viable cellular area,

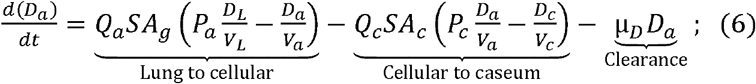

and drug amount (in mg) within caseum,

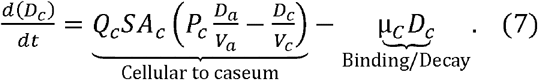

We update the predicted volumes of these regions *V*_*a*_, *V*_*c*_ and the caseum surface area *SA*_*C*_ hourly based on cell numbers within a virtual granuloma and macrophage death that contributes to caseum, respectively (see Appendix A). The drug-penetration coefficients *Q*_*a*,_ *Q*_*c*,_ *P*_*a*,_ *P*_*c*_ principally drive antibiotic levels within the cell area and caseum. The parameters μ_*C*_ and μ_*D*_ represent decay rate constants in the caseum and the viable cellular area, although we assume that the decay rate constant in uninvolved lung tissue and the viable cellular region of the granuloma is identical. It is important to note that *D*_*c*_ represents how much drug has permeated into the caseated volume, which should not be confused with the drug’s ability to uniformly distribute within the caseum (see 4.5.2).

For each drug and each host, we generate an independently-parameterized copy of the equations for 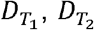 (where applicable), *D*_*PT*,_ *D*_*B*_ and *D*_*L*_ For each drug and granuloma pair, we independently parameterize copies of the equations guiding *D*_*a*_ and *D*_*c*_ We calibrate parameters of all PK equations using drug concentration datasets from various tissue types (e.g., blood, caseum, uninvolved lung, cellular or necrotic lesions) in humans^39,84^ or rabbits^36,85^. We calculate PK during multi-drug regimen simulations assuming that drug-drug interactions do not affect PK. We present calibrated parameters in **Appendix A**.

#### 4.5.2 Submodel of drug penetration into caseum

Within a granuloma, nonreplicating bacteria *B*_*N*_ may be distributed throughout caseum. However, using a single compartment for antibiotics within caseum (*D*_*C*_) implicitly assumes that the concentration is equally distributed throughout the entire caseum (well-mixed). For some drugs, this is not far from the truth (e.g. INH and PZA), although many drugs bind to macromolecules in caseum and fail to penetrate (e.g., BDQ)^55^.

Within *GranSim*, which follows just a single granuloma, we track the spatial distribution of cells within a granuloma, and thus we can capture spatial heterogeneity. However, for computational feasibility in *HostSim*, which follows multiple granulomas within a single host, we represent cellular and caseum regions as two physiological compartments comprising a granuloma. Thus, we assume a worst-case scenario wherein all non-replicating bacteria *B*_*N*_ reside at the center of a spherical volume of caseum of radius *r*_*C*_ with a reduced ability for drugs to penetrate. We assume this is based on a fractional binding by caseum macromolecules. We represent this by calculating PD for *B*_*N*_ killing with 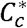 for each drug, where

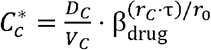

Here, *r*_0_ = 0.4 mm is the median diameter of a granuloma in *HostSim*, and β is the effective fraction of *C*_*C*_ buried within *r*_0_ of caseum. The relative concentration of drugs just inside of caseum and in the center of caseum has been quantitatively and spatially mapped in experimental PK studies^55^. For all drugs, we apply the fitting parameter τ = 0.273 to adjust 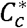 such that we have ≈58 %reduction of MXF and ≈ 80 %of RIF at the center of 0.2mm of caseum based on those results. The value of β defines a distribution-like function relating the drug’s affinity to caseum to the size of the granuloma. To ensure accurate relative caseum-binding affinities, we chose values of β to be equal to the fractional unbound percentages of drug estimated in experiments^55^. We used these penetrating concentrations when computing PD of each drug, discussed in detail in the next subsections.

#### 4.5.3 Pharmacodynamics (PD) modeling

Once we compute concentrations of each active drug within each granuloma compartment (and penetrating concentrations within caseum), we represent the drugs’ activity by a Hill equation adapted from previous *GranSim* work^24,25^:

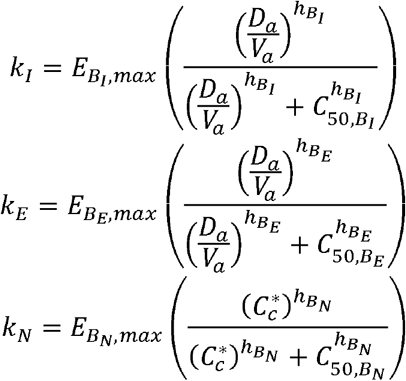

where *k*, the killing rate constant is a function of the drug concentration, *E*_max_ is the maximum killing rate, *C*_5o_ is the concentration needed to achieve half maximum drug action (*E*_max_ /2) and *h* is the Hill constant. Mtb are in distinct metabolic states in different locations (intracellular, extracellular or trapped in caseum) and they metabolize drugs differently in each. We represent drug action separately for Mtb in each location: nonreplicating Mtb *B*_*N*_ trapped in caseum, intracellular Mtb *B*_*I*_ within macrophages and extracellular replicating Mtb *B*_*E*_ in the granuloma cellular area. For single-drug regimens, we use this kill rate constant within an additional death term for Mtb populations:

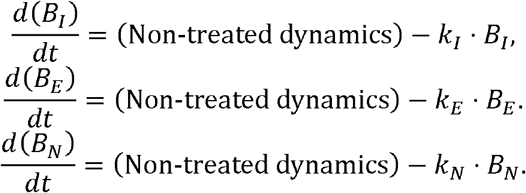

For multiple-drug regimens, we compute interactions as described in the next subsection. We calibrate Hill curve parameters *E*_max,_ *C*_5o_ and *h* using *in vitro* bactericidal assays. Namely, we use assays in caseum mimic, macrophage assays and Mtb assays to calibrate parameters for nonreplicating, intracellular and extracellular Mtb, respectively (**Table 3**). We used these calibrated parameters in our previous *GranSim* work^25,40^.

#### 4.5.2 Minimum therapeutic concentrations for pharmacodynamics

Antibiotics are not effective unless they are present in sufficiently high concentrations. For this reason, we distinguish each antibiotic as being *active* in a compartment only if it exceeds experimentally-observed minimal concentrations (see **Table 4**). In all PD and multi-drug interaction sections below, we refer only to those drugs that are active in all computations. If all drugs are below-threshold, we consider the treatment as ineffective, and all kill-rates are set to 0.

**Table 4:**
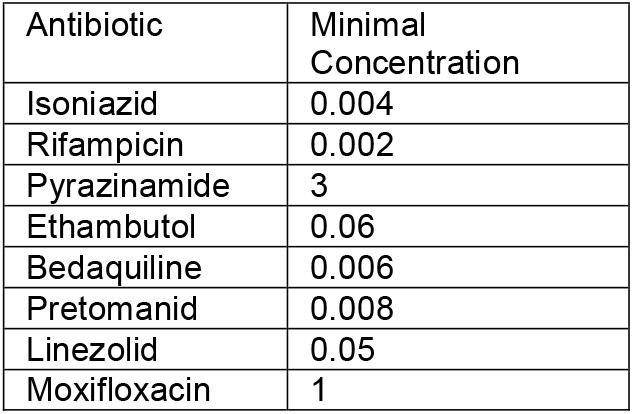
Minimal concentrations (in mg/L) used to determine whether drugs in *HostSim* are considered active. All values taken from literature^91^.

|  |  |
| --- | --- |
| Antibiotic | Minimal Concentration |
| Isoniazid | 0.004 |
| Rifampicin | 0.002 |
| Pyrazinamide | 3 |
| Ethambutol | 0.06 |
| Bedaquiline | 0.006 |
| Pretomanid | 0.008 |
| Linezolid | 0.05 |
| Moxifloxacin | 1 |

To illustrate this, we present a simple example. If we administer HRZM to a virtual host and the INH concentration is below-threshold in the viable cellular area of one granuloma, then we compute PD values within the viable cellular area as though we were treating the host with RZM. If, within the caseum of that same granuloma at the same time, we calculate that INH and MXF have effective concentrations (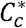 values) below the minimal threshold,then we calculate PD as though we were treating the host with RZ.

#### 4.5.5 Modeling drug-drug interactions when used in combination regimens

When drugs are used in combination, they can interact with each other resulting in synergistic or antagonistic effects. We incorporate drug-drug interactions when multiple drugs are present within a physiological compartment (e.g., cellular area or caseum). Briefly, we adjust effective concentrations of active drugs using fractional inhibitory concentrations (FICs) of drug combinations predicted by an *in silico* tool, INDIGO-MTB (inferring drug interactions using chemogenomics and orthology optimized for Mtb)^83,98^, as we have done previously^25,40,59^. INDIGO-MTB is a machine learning based tool that uses known drug interactions and drug transcriptomics data to predict unknown drug interactions in the form of FICs. FIC values lower or higher than 1 mean the drugs are synergistic or antagonistic, respectively, whereas an FIC value of 1 means the drugs do not interact, i.e., they are additive.

We model drug interaction by converting concentrations of each active drug *i*(*C*_*i*_) to equipotent concentrations of drug *i*_*max*_ the drug with highest maximal killing rate (i.e., highest *E*_*max*_) To do that, we calculate the adjusted concentration of *i*(*C*_*i,adj*_) which is the concentration of *i*_*max*_ that would kill Mtb with the same rate as drug *i* with concentration *C*_*i*_.

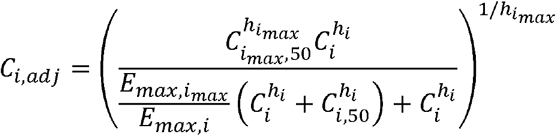

where 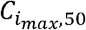 and *C*_*I*,50_ are the concentration of *i*_*max*_ and *i* at which half maximal killing is achieved, respectively, 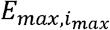 and *E*_*max*,*i*_ are the maximal killing rate constants of drug *i*_*max*_ and drug I respectively, and 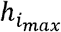 and *h*_*i*_ are the Hill coefficients of drug *i*_*max*_ and drug *i* respectively. Once we calculate the adjusted concentrations of all drugs in a compartment, we determine the effective concentration (*C*_*eff*_) from:

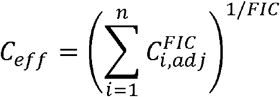

where *C*_*i*,*adj*_ is the adjusted concentration of drug *i*,*n* is the number of drugs in a compartment and *FIC* is the *FIC* value predicted for these *n* drugs by INDIGO-MTB. Then, we calculate the effective killing rate constant *k* by using *C*_*eff*_ and Hill parameters of 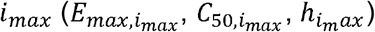:

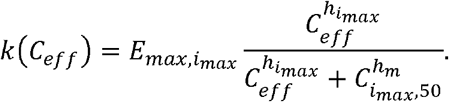

#### 4.5.6 Drug sterilization of bacteria trapped within infected macrophages (B_I_)

In *HostSim*, we pool the entire population of intracellular bacteria *B*_*I*_ within each granuloma across all macrophages. If there are many bacteria per infected macrophage, it stands to reason that killing a single intracellular bacterium does not clear any individual macrophage. By contrast, antibiotic killing of many bacteria is likely to reduce the number of infected macrophages. The exact collection of parameters that control the portion of infected macrophages that clear their infection is unclear, but we assume that there are several factors that are driven by granuloma-driven (i.e., spatial arrangement of granuloma, heterogeneity of macrophages and Mtb) while others that are driven by the PK of individual drugs. Moreover, it has been known for some time that some antibiotics have higher intracellular/extracellular concentration ratio (*R*_*IC* / *IC*_) in alveolar macrophages (see **Table 5**) than others^99,100^. We assume that both factors are relevant in determining overall sterilization of bacteria within macrophages.

**Table 5:** Measured ranges of the ratio of intracellular drug concentration to extracellular drug concentration from literature. We used the mean ratio in *HostSim* to determine rates of macrophage clearance.

| Antibiotic | Minimum concentration ratio | Maximum concentration ratio | Source | Mean $R_{IC/EC}$ |
| --- | --- | --- | --- | --- |
| Isoniazid | 0.8 | 1.8 | <sup>39</sup> | 1.3 |
| Rifampicin | 4.4 | 8.6 | <sup>39</sup> | 6.5 |
| Pyrazinamide | 0.5 | 0.7 | <sup>39</sup> | 0.6 |
| Ethambutol | 1.8 | 18.8 | <sup>99,101</sup> | 10.3 |
| Bedaquiline | 47 | 917 | <sup>102</sup> | 482 |
| Pretomanid | 4.6 | 8.6 | <sup>91</sup> | 6.6 |
| Linezolid | 1.3 | 5.9 | <sup>102,103</sup> | 3.6 |
| Moxifloxacin | 15 | 20 | <sup>104</sup> (see also <sup>105</sup>) | 17.5 |

**Table 6:** Regimens studied in previous work and recreated herein with *HostSim*. Each column indicates a concentration, in mg/kg to be administered for 6 months (180 days). (*) This concentration is given only for one month, while other drugs continue to be administered. (†) Note that in each regimen set, we use the nomenclature from the original study; HRZE in set (iii) has different dosages from (i) and (ii).

| Regimen Name | Days/week | INH | RIF | PZA | EMB | BDQ | PTM | LZD | MXF |
| --- | --- | --- | --- | --- | --- | --- | --- | --- | --- |
| Regimen set (i) - HRZE versus BDQ and MXF-containing regimens. |  |  |  |  |  |  |  |  |  |
| HRZE | 7 | 5 | 10 | 25 | 20 | - | - | - | - |
| RMZE | 7 | - | 10 | 25 | 20 | - | - | - | 7 |
| BPaL | 7 | - | - | - | - | 20 | 20 | 90 | - |
| BPa | 7 | - | - | - | - | 20 | 20 | - | - |
| HRZM | 7 | 5 | 10 | 25 | - | - | - | - | 7 |
| BPaZ | 7 | - | - | 25 | - | 20 | 20 | - | - |
| BPaLZ | 7 | - | - | 25 | - | 20 | 20 | 90 | - |
| BPaM | 7 | - | - | - | - | 20 | 20 | - | 7 |
| BPaMZ | 7 | - | - | 25 | - | 20 | 20 | - | 7 |
| PaMZ | 7 | - | - | 25 | - | - | 20 | - | 7 |
| BPaLM | 7 | - | - | - | - | 20 | 20 | 90 | 7 |
| BPaL-1moL | 7 | - | - | - | - | 20 | 20 | 90* | - |
| Regimen set (ii) - Marmoset studies from *GranSim* study<sup>40</sup>. |  |  |  |  |  |  |  |  |  |
| RMZE | 7 | - | 10 | 25 | 20 | - | - | - | 7 |
| BPa | 7 | - | - | - | - | 20 | 20 | - | - |
| BPaL | 7 | - | - | - | - | 20 | 20 | 90 | - |
| BL | 7 | - | - | - | - | 20 | - | 90 | - |
| RM | 7 | - | 10 | - | - | - | - | - | 7 |
| HRZE | 7 | 5 | 10 | 25 | 20 | - | - | - | - |
| PaL | 7 | - | - | - | - | - | 20 | 90 | - |
| Bedaquiline | 7 | - | - | - | - | 20 | - | - | - |
| Pretomanid | 7 | - | - | - | - | - | 20 | - | - |
| RZ | 7 | - | 10 | 25 | - | - | - | - | - |
| HZ | 7 | 5 | - | 25 | - | - | - | - | - |
| Moxifloxacin | 7 | - | - | - | - | - | - | - | 7 |
| Pyrazinamide | 7 | - | - | 25 | - | - | - | - | - |
| Rifampicin | 7 | - | 10 | - | - | - | - | - | - |
| Isoniazid | 7 | 5 | - | - | - | - | - | - | - |
| Regimen set (iii) - Clinically ranked and *GranSim*-ranked regimens<sup>50,59</sup>. |  |  |  |  |  |  |  |  |  |
| H10R10E25 | 7 | 10 | 10 | - | - | - | - | - | - |
| H10E25 | 7 | 10 | - | - | 25 | - | - | - | - |
| H6E15 | 7 | 6 | - | - | 15 | - | - | - | - |
| H6E25 | 7 | 6 | - | - | 25 | - | - | - | - |
| H6E6 | 7 | 6 | - | - | 6 | - | - | - | - |
| HR | 7 | 5 | 11 | - | - | - | - | - | - |
| H6R10E16 | 7 | 6 | 10 | - | 16 | - | - | - | - |
| H6R6E10 | 7 | 6 | 6 | - | 10 | - | - | - | - |
| R11E25 | 7 | - | 11 | - | 25 | - | - | - | - |
| H11R10 | 7 | 11 | 10 | - | - | - | - | - | - |
| H15R10E40dpw2 | 2 | 15 | 10 | - | 40 | - | - | - | - |
| H6R10E25 | 7 | 6 | 10 | - | 25 | - | - | - | - |
| H6R9Z24E16dpw3 | 3 | 6 | 9 | 24 | 16 | - | - | - | - |
| R23.5E45dpw2 | 2 | - | 23.5 | - | 45 | - | - | - | - |
| R23.5E90dpw1 | 1 | - | 23.5 | - | 90 | - | - | - | - |
| R24Z64E90dpw1 | 1 | - | 24 | 64 | 90 | - | - | - | - |
| R23.5E25 | 7 | - | 23.5 | - | 25 | - | - | - | - |
| H5R10Z25E17.5 | 7 | 5 | 10 | 25 | 17.5 | - | - | - | - |
| H14R13Z47E29dpw3 | 3 | 14 | 13 | 47 | 29 | - | - | - | - |
| H16R12Z49dpw2 | 2 | 16 | 12 | 49 | - | - | - | - | - |
| HRZ | 7 | 5 | 10 | 25 | - | - | - | - | - |
| H5R10M7Z25dpw5 | 5 | 5 | 10 | 25 | - | - | - | - | 7 |
| H14R10M9Z34dpw3 | 3 | 14 | 10 | 34 | - | - | - | - | 9 |
| H6R10Z27E16 | 7 | 6 | 10 | 27 | 16 | - | - | - | - |
| HRZE† | 7 | 5 | 10 | 33 | 25 | - | - | - | - |
| R10M7Z25E15 | 7 | - | 10 | 25 | 15 | - | - | - | 7 |

To capture both host factors and drug factors, we add the following terms to macrophage and caseum dynamics within *HostSim*.

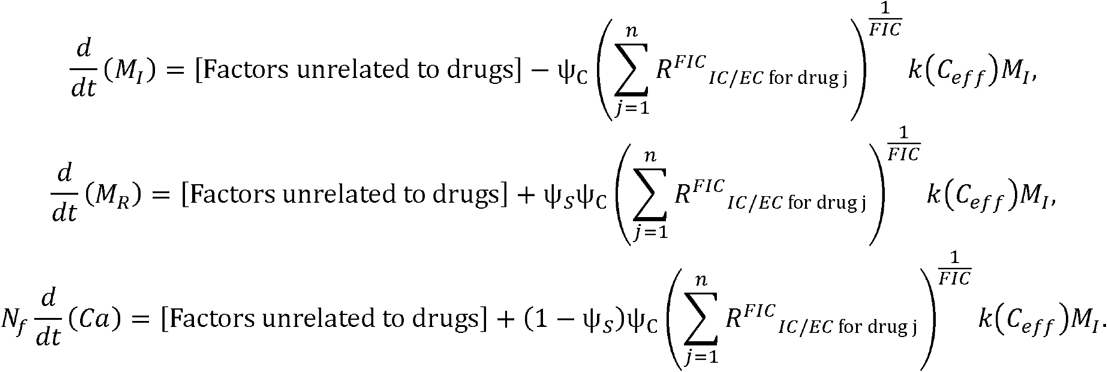

For the complete equations for *M*_*I*,_ *M*_*R* and_ *C*_*a*_, see Section 1 of **Appendix A**.

Here, Ψ_c_ is a fitting parameter encompassing the granuloma-driven factors of macrophage clearance and is sampled and considered as a granuloma-scale parameter. The variable *n* is the number of active drugs, and we use the values from **Table 5** for a drug-interaction-informed ratio of intracellular to extracellular drug concentration. After the intracellular bacteria within macrophages are cleared, we assume that macrophages either revert to a resting state or are killed; this outcome is controlled by a survival parameter (Ψ_c_).

### 4.6 Simulated multi-antibiotic regimens

Nearly every approved treatment for TB involves regimens of multiple antibiotics. Accordingly, we must represent multiple drugs in *HostSim*—including the standard CDC regimen of HRZE. We also choose several new and repurposed drugs (B, Pa, L, M), as they have been shown to potentially improve treatment for Mtb infection based on preclinical and clinical trials^47,49^.

Our previous work^40,59^ investigated granuloma-scale drug-ranking in-depth using *GranSim*. There, we investigated several datasets at the granuloma-scale that we now represent at the whole-host scale (**Table 6**): (i) HRZE versus BDQ, MXF, and PTM-containing regimens that are promising to shorten treatment time based on clinical and preclinical trials^41-43,45-49,51,61,65,66,106-114^; (ii) a series of antibiotic regimens used in marmoset studies^40^; and (iii) a set of 26 drug regimens that were clinically ranked in a previous meta-analysis^50^ and further explored by *GranSim*^*59*^.

### 4.7 Quantifying the efficacy of virtual host response to drugs

To validate our model, we compare measurements taken from experimental or clinical contexts with analogous simulation outputs. Note that all granuloma-scale outcomes are analyzed only by examining primary (i.e., non-disseminated) granulomas to control for time-post-granuloma-formation. We also acknowledge that we are modeling a disease process that occurs in humans, NHPs, and other biological model systems; accordingly, we detail how each dataset is comparable to *HostSim* and by which measure below.

#### 4.7.1 Early bactericidal activity (EBA) studies

EBA assays have been used for decades to determine effectiveness of TB antibiotic treatments on reducing sputum bacillus count within the first two weeks of treatment^115^. Although these studies are limited to studying bacteria that are present in sputum, they represent a noninvasive measurement of efficacy early during drug development measured in most anti-TB antibiotics.

We virtually replicate a measurement of EBA by calculating the mean rate of change of log_10_ CFU in sputum samples between start-of-treatment up to two weeks later^116.^ We calculate our virtual host EBA value as:

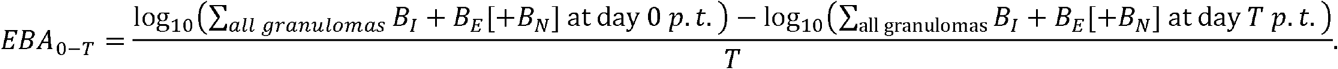

It is unclear whether any bacteria trapped in caseum (*B*_*N*_) will contribute to such sputum measurements, as this pertains to the rate and frequency with which *B*_*N*_ may be removed from caseum. We measure EBA both with and without *B*_*N*_ contribution, described in the results text as relevant. Interestingly, mono-INH treatment is used as a positive-control as it reports a very high *EBA*_0 −2_ compared to other drugs, albeit with high variance and compared to other drugs, albeit with high variance and despite INH having relatively low efficacy compared to other antibiotics^115^. This may be due to a partial contribution of non-replicating bacteria to the counts present in sputum, and INH having a relatively high killing rate of extracellular Mtb.

#### 4.7.2 Sterilization versus Apparent-sterilization of virtual hosts

Historically, the gold standard for determining if a patient had sterilized Mtb infection was to perform a solid culture on Löwenstein-Jensen (LJ) media. Although this is no longer the preferred method of diagnosis due to the longer incubation times than liquid media such as MGIT 960^117,118^, solid LJ culture was used in many Phase IIb studies and a 2021 meta-analysis used this measurement to rank treatment regimens^50^. We have previously simulated this measurement for validation purposes in *GranSim*^*59*^. Culture conversion is not a precise measurement; small numbers of viable bacilli may not result in culture conversion. To capture the impact of a limit of detection in *HostSim*, we define a virtual host as *apparently-sterilizing* if there are less than 50 total CFU in the virtual host and a virtual granuloma as *apparently-sterilizing* if it has fewer than 10 CFU. This is used in virtual host classification and ranking, below.

#### 4.7.3 Classification of Virtual Hosts

In *HostSim*, we have two heuristics that we use to classify a virtual host as having active TB disease. First, the virtual host classified as having active TB if it maintains more than a total of CFU for more than 30 days; the CFU count was experimentally observed as being correlated to progressive infection^26^. Second, the virtual host has active TB disease if the CFU count of any of its granulomas increases by a factor of 1.25 between days 70 and 365 post-infection, indicating continued long-term growth. We allow the behavior of one granuloma to dictate the host state because this is known to be true of TB^78^. To understand the role of capturing low-CFU infections, we distinguish between hosts that are *sterilizing hosts* (total CFU = 0) and *apparently-sterilizing* (total lung CFU < 50; note our model is flexible to vary this number). We classify all other hosts as having subclinical infection. In some contexts, we compare hosts with subclinical infection to apparently-sterilizing hosts and in other contexts we compare hosts with subclinical infection to sterile hosts. In each context, classifications of hosts always create disjoint sets of hosts—i.e., when examining apparent-sterilization, a host with 7 CFU will be classified as apparently-sterilizing instead of as subclinically-infected. When examining sterilization of populations, this same 7-CFU host will be considered as having subclinical infection. This distinction is crucial in considering the importance of Mtb detection thresholds in drawing whole-population conclusions.

### 4.8 Drug regimen rankings

A key goal in many studies of antibiotics is to rank efficacy of antibiotic regimens. Methods of ranking all are developed towards an intuition of “goodness-of-drug”, or even a more concrete feature such as “sterilizing potential”. Regardless, multiple considerations must be considered in constructing a ranking, including: (i) data availability, (ii) detectability of bacteria in small numbers, (iii) initial bacterial burden of hosts in the trial cohort, and (iv) scale of the ranked data.

Our simulated data contains a single virtual cohort of 500 virtual hosts, each of having at least one non-sterilizing granuloma prior to treatment (300 days p.i.). We perform each ranking at both the granuloma-scale and host-scale (e.g., using individual granuloma CFU or total lung CFU, respectively). We also perform rankings for either (i) all hosts in the virtual cohort, (ii) high-CFU hosts only, or (iii) low-CFU hosts only. We determine a granuloma as high-CFU if it has >1,000 CFU prior to treatment, and a host as high-CFU if it has >10,000 total CFU prior to treatment. Finally, we define a new ranking method based relative to *sterilization* of hosts/granulomas (<0.5 CFU), or *apparent-sterilization* of hosts/granulomas (<50 and <10 CFU, respectively).

Our rankings are based on all virtual hosts and their primary granulomas, as well as granulomas that disseminate before the start of treatment (as treatment may affect dissemination probability). At the host scale, 496 virtual hosts do not sterilize prior to infection, of which 53 are high-CFU and 443 are low-CFU. At the granuloma scale, we consider 5729 non-sterile granulomas (pooled from across all hosts), of which 5413 are low-CFU and 316 are high-CFU. Without loss of generality, we describe our ranking methods in terms of measuring a single population (e.g., in terms of sterilization of virtual granulomas), but these methods may be modified for the different subpopulations (e.g., we may rank based on apparent-sterilization of high-CFU hosts).

#### 4.8.1 Area under sterilization curve (AUSC)-based ranking

This method incorporates the sterilization history of the virtual population, rather than just the portion of sterilized hosts/granulomas at one time point. We have previously developed this method for ranking regimens within a regimen set using *GranSim*^40^.

First, we randomly assign each granuloma into one of five equal-sized groups (four groups of 1146 and one group of 1145). Then, for each group, we record the fraction of those granulomas that sterilize at each time to generate a sterilization curve for each group. Then, we calculate the area under the sterilization curve (A USC) for each group’s sterilization curve, resulting in a sample of five AUSC values for each regimen *S*(*R*). We assign a regimen *R*_*i*_ is assigned if *S*(*R*_*i*_) has a statistically significantly larger mean AUSC value than another regimen *S*(*R*_*j*_) and we penalize the regimen a point if it has a statistically significantly lower mean AUSC value than another regimen. That is, we define δ(*i*,*j*) =1 if *S*(*R*_*i*_) is significantly larger than *S*(*R*_*j*_) and 0 otherwise;

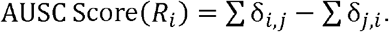

We then rank each regimen based on their scores, with a rank of 1 being the highest score. If *n* regimens share a score, we assign them the same rank, and the ranking of regimen with the next-lowest score is *n* lower. As an example, if regimens A, B, C, D have AUSC scores 2, 2, -1, -3, then we assign their AUSC rankings as 1, 1, 3, and 4, respectively.

#### 4.8.2 Ranking based on simulated 8-week solid-culture tests

This measurement is an *in-silico* emulation of a solid-culture test (see Section 4.7.2). In this ranking method, we examine the percentage of our virtual hosts that are apparently-sterilizing (<50 total CFU) at eight weeks post-treatment-start. We assign better (lower) ranks to regimens with higher percentages, with the best-sterilizing regimens being ranked as rank 1. We address tied ranks as we do with the AUSC method.

### 4.9 Multi-scale intervention analysis of virtual host-drug response

A central advantage of computational models is that we are able to run and re-run many virtual experiments <u>using the same virtual subjects under different conditions</u>. Measurements such as EBA compare some early time-points to later time-points to understand the action of drugs within individuals (Section 4.7). By using simulations, we are able to examine the response of one virtual host (in the sense of one virtual identity, Section 4.3) represented in different *what-if* scenarios. In this way, we can generate detailed information about host-scale heterogeneity of intervention efficacy by using a multi-scale interventional design (MID) framework for identifying features of virtual TB hosts and granulomas that correlate with their drug response^26^.

Briefly, MID works by defining a collection of virtual hosts, and then using two different model scenarios— control and intervention—to generate each virtual host outcome in either scenario. We can then score each host’s level improvement at every time point in the simulation by comparing the virtual host outcome in the with-treatment / without-treatment scenarios. Doing this requires defining an *impact score* function that compares the control and intervention scenario outcomes per-timepoint. We then use impact score as a model output dependent only on virtual host identity so that we can then examine factors driving host-response heterogeneity by using sensitivity analysis methods such as PRCC (see Section 4.10).

#### 4.9.1 Multi-scale interventional design scores

We rigorously define notions of host improvement, as different scoring methods may result in different conclusions drawn about intervention efficacy. For instance, a trial may report a drug with a high EBA but that has poor sterilization of persister populations. Although it may be reasonable to assume that “fewer CFU is better for a patient”, assigning scores to partially-reduced bacterial burden bears consideration. To cover several definitions of improvement, we define multiple impact scores measuring host improvement that may seem semantically similar: (1) total CFU reduction, (2) non-replicating CFU reduction, (3) percent reduction of total CFU, and (4) percent reduction of non-replicating CFU. The first two scores look at CFU differences between the untreated virtual hosts/granulomas and how CFU would evolve during treatment:

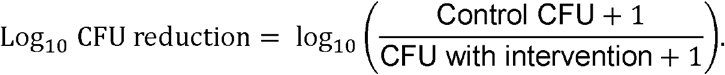

Similarly, we examine reduction of non-replicating bacteria:

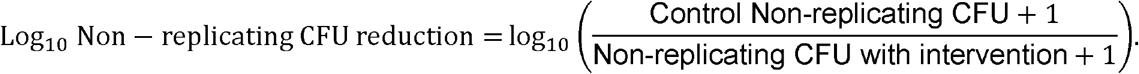

As a remark, if we assume that a virtual host in the control (no treatment) scenario has a relatively constant bacterial burden over two weeks, then dividing total CFU reduction by time-since-treatment-start is similar to EBA (Section 4.7), i.e.*EBA*_*0* −*T*_ ≈(Log_10_ CFU reduction)/*T*

The final two scores, percent CFU reduction, capture the portion of CFU burden (total or non-replicating) that each drug-treated granuloma has relative to what it would-have-had without treatment.

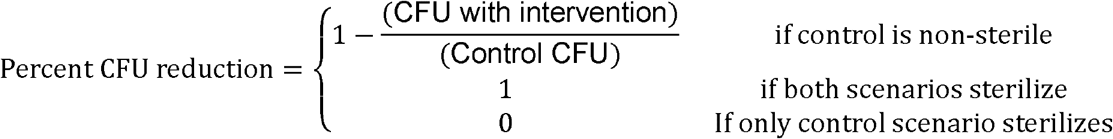

and

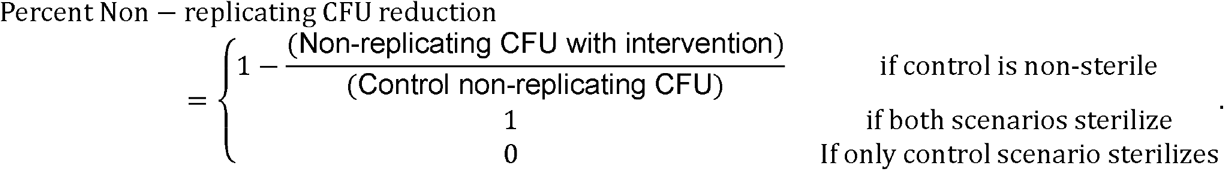

In each case, we can evaluate these scores at any scale. At the granuloma scale, “Control CFU” refers to the CFU within a granuloma. Similarly, at the host scale, “Control CFU” refers to the total CFU from all granulomas within a virtual host.

### 4.10 Sensitivity analysis identifies major drivers of outcomes

Biologically, there are vast differences between people based on differences in their anatomy, metabolism and physiology, and these differences occur within ranges that still result in an identifiably healthy individual. Similarly, the differences between virtual host identities in our *HostSim* virtual cohort come in the form of distinct parameter choices - the specific numerical values that determine the rates and relative activity of different mechanisms (e.g., cytokine secretion rates, saturation points, carrying capacities, etc.). A key goal of *in silico* experiments is to determine the relative importance of varying individual parameters, thereby identifying which mechanisms are principally responsible for variations within simulated outcomes. To do this, we use uncertainty and sensitivity analyses, which encompasses many well-developed methods used to determine how parameter uncertainty affects model output(s)^119-124^. Global sensitivity analysis is an approach that looks at variations across all of the parameter space^125^—i.e., how every parameter’s variation affects the outcomes while considering nonlinear interactions. By contrast, global sensitivity analysis methods (e.g., Sobol’ index method^126-128^, partial rank correlation coefficient (PRCC) method^76,129-131^, or eFAST^76,120^) function by evaluating the model at many points in parameter and consequently are not suitable for extremely computationally intensive models, as hundreds or thousands of model evaluations must be performed. In *HostSim*, our large number of variable parameters and fast model wall-time makes it an ideal use-case for global sensitivity analysis.

We use the PRCC method^76^ to quantify the relative contribution of parameters to simulated heterogeneity. This method generates, for each parameter, a nonlinear correlation value (a PRCC value) between model parameters or initial conditions and a given model outcome. PRCC may also be applied to outputs that compare any two models, as long as those models depend on the same parameter collections (see Section 4.9). This method also determines statistical significance of multiple PRCC values, Benjamini-Hochberg corrected for multiple-comparisons.

Recall that *HostSim* generates separate ODE systems for each granuloma within a host but parameterizes them such that granulomas within a host are more likely to be similar to those between hosts (Section 4.3, **Figure 7**A). Most sensitivity analysis methods assume that models are either a single in/out black-box or, as in hierarchical methods, parameter ranges are not dependent upon larger-scale components^125^. Using the multi-scale structure of *HostSim*, we can obtain PRCC values separately at each scale, i.e. a *multi-scale sensitivity analysis*. At the whole-host scale, we consider host baseline parameter values as inputs that generate whole-host outcomes—e.g., total host CFU. We then analyze the same mechanisms at the granuloma scale by comparing parameters against same-scale outcomes (e.g., granuloma CFU). In this way, we can determine (i) which mechanisms impact outcomes at the smallest scale, and (ii) whether host baselines are sufficient to predict host outcomes (**Figure 7**B).

### 4.11 Methods for new for non-human primate data

#### Bedaquiline treatment study in macaques

Five adult male Mauritian cynomolgus macaques were obtained from Bioculture, Inc (Mauritius), with an age range of 5-9 years. Macaques were co-housed in a Biosafety Level 3 facility and infected with 22 CFU M. tuberculosis strain Erdman via bronchoscope as described previously^132^. Macaques were monitored clinically throughout the course of the experiment. Eight weeks after M. tuberculosis infection, the animals were treated with the BPaL regimen daily via oral administration in food treats for 4 weeks. Drug doses: Bedaquiline 40mg/kg, Pretomanid 62 mg/kg, Linezolid 20 mg/kg. Each lung granuloma sample was plated for bacterial burden individually on 7H10 plates; the plates were incubated at 37 degrees with 5% CO2 and counted for colony forming units (CFU) after 3 weeks. Total lung burden was determined by summing all samples from lungs. The fraction (%) of sterile (CFU negative) granulomas was calculated for each animal.

## Appendix A

HostSim *technical description. This document contains all of the* HostSim *equations, parameter ranges, and a description of how the different components are linked together. This document is available on-line at http://malthus.micro.med.umich.edu/lab/supplements/HostSim-3/* .

## Acknowledgements

The authors thank Dr. Sriram Chandrasekaran for fractional inhibitory concentration (FIC) values obtained from his Inferring drug Interactions using chemo-genomics and orthology (INDIGO) studies. They thank Dr. George Drusano for detailed access to pretomanid bactericidal assays. We thank Paul Wolberg for computational assistance and support. This work was funded through the Gates Medical Research Institute (D.K.) and by National Institutes of Health Grant R01 AI50684 (D.K.) and the Center for Data-Driven Drug Development and Treatment Assessment (DATA; D.K. and M.B.). C.T.M. was supported by the Molecular Mechanisms in Microbial Pathogenesis Training Program (T32 AI007528).

## Notes

http://malthus.micro.med.umich.edu/lab/supplements/HostSim-3/

### Competing Interest Statement

The authors have declared no competing interest.

### Summary of Updates

Revised text of the abstract. Corrected author information in the manuscript PDF.

