## Supplemental Material 1 for "Rankings of tuberculosis antibiotic treatment regimens are sensitive to spatial scale, detection limit, and initial host bacterial burden"

Christian T. Michael[1†], Maral Budak[1†], Pauline Maiello[2], Kara Kracinovsky[2], Mark Rodgers[2], Jaime Tomko[2], Philana Ling Lin[3], JoAnne Flynn[2], Jennifer J. Linderman[4], Denise Kirschner[1]
[1]Department of Microbiology & Immunology, University of Michigan - Michigan Medicine, Ann Arbor, MI, USA
[2]Microbiology and Molecular Genetics, University of Pittsburgh School of Medicine, Pittsburgh, PA, USA
[3]Department of Pediatrics, Children's Hospital of the University of Pittsburgh of UPMC, Pittsburgh, PA, USA
[4]Department of Chemical Engineering, College of Engineering, University of Michigan, Ann Arbor, MI, USA

### 1 Supplemental figures and tables

**Supplemental Table 1: Parameters that are significantly correlated to efficacies of HRZE, RMZE and BPaL as measured by $log_{10}$ non-replicating CFU reduction in granuloma and host scales for at least 5 consecutive days (alpha = 0.05).** First column and second column refer to the first 50 days and after 50 days of treatment, respectively. ++ and –- indicate that the PRCC value is 0.4 to 0.6 and -0.4 to -0.6, respectively. + and - indicate that the PRCC value is 0.2 to 0.4 and -0.2 to -0.4, respectively.

<table>
<tr><th>Parameter Description</th><th colspan="2">Correlation with HRZE efficacy, Host-scale, Early | Late</th><th colspan="2">Correlation with HRZE efficacy, Granuloma-scale, Early | Late</th><th colspan="2">Correlation with RMZE efficacy Host-scale, Early | Late</th><th colspan="2">Correlation with RMZE efficacy Granuloma-scale, Early | Late</th><th colspan="2">Correlation with BPaL efficacy Host-scale, Early | Late</th><th colspan="2">Correlation with BPaL efficacy Granuloma-scale, Early | Late</th></tr>
<tr><td>Fraction of macrophage-released bacteria that get trapped in caseum (CN)</td><td>+</td><td>++</td><td>++</td><td>++</td><td>++</td><td>++</td><td>++</td><td>++</td><td>++</td><td>++</td><td>++</td><td>++</td></tr>
<tr><td>Growth rate constant of extracellular bacteria (alpha20)</td><td>++</td><td>++</td><td>+</td><td>++</td><td>++</td><td>++</td><td>++</td><td>++</td><td>++</td><td>++</td><td>++</td><td>++</td></tr>
<tr><td>The rate constant of nonreplicating bacteria in caseum reverting to a replicating state (krev)</td><td></td><td>-</td><td></td><td>-</td><td>-</td><td>-</td><td></td><td>-</td><td></td><td>-</td><td></td><td>-</td></tr>
<tr><td>Rate constant of TNF-driven recruitment of resting macrophages to granulomas (Sr4b)</td><td></td><td></td><td></td><td></td><td></td><td></td><td></td><td></td><td>-</td><td></td><td></td><td></td></tr>
</table>

**Supplemental Table 2: Parameters that are significantly correlated to efficacies of HRZE, RMZE and BPaL as measured by percent CFU reduction in granuloma and host scales for at least 5 consecutive days (alpha = 0.05).** First column and second column refer to the first 50 days and after 50 days of treatment, respectively. ++ and –- indicate that the PRCC value is 0.4 to 0.6 and -0.4 to -0.6, respectively. + and - indicate that the PRCC value is 0.2 to 0.4 and -0.2 to -0.4, respectively.

<table>
<tr><th>Parameter Description</th><th colspan="2">Correlation with HRZE efficacy, Host-scale, Early | Late</th><th colspan="2">Correlation with HRZE efficacy, Granuloma-scale, Early | Late</th><th colspan="2">Correlation with RMZE efficacy Host-scale, Early | Late</th><th colspan="2">Correlation with RMZE efficacy Granuloma-scale, Early | Late</th><th colspan="2">Correlation with BPaL efficacy Host-scale, Early | Late</th><th colspan="2">Correlation with BPaL efficacy Granuloma-scale, Early | Late</th></tr>
<tr><td>Fraction of macrophage-released bacteria that get trapped in caseum (CN)</td><td></td><td></td><td>-</td><td></td><td></td><td></td><td></td><td></td><td></td><td></td><td>-</td><td></td></tr>
<tr><td>Growth rate constant of extracellular replicating bacteria (alpha20)</td><td>-</td><td></td><td>-</td><td></td><td></td><td></td><td></td><td></td><td>-</td><td></td><td>-</td><td></td></tr>
</table>

**Supplemental Table 3: Parameters that are significantly correlated to efficacies of HRZE, RMZE and BPaL as measured by percent non-replicating CFU reduction in granuloma and host scales for at least 5 consecutive days (alpha = 0.05).** First column and second column refer to the first 50 days and after 50 days of treatment, respectively. ++ and –– indicate that the PRCC value is 0.4 to 0.6 and -0.4 to -0.6, respectively. + and - indicate that the PRCC value is 0.2 to 0.4 and -0.2 to -0.4, respectively.

| Parameter Description | Correlation with HRZE efficacy, Host-scale, Early \| Late | | Correlation with HRZE efficacy, Granuloma-scale, Early \| Late | | Correlation with RMZE efficacy Host-scale, Early \| Late | | Correlation with RMZE efficacy Granuloma-scale, Early \| Late | | Correlation with BPaL efficacy Host-scale, Early \| Late | | Correlation with BPaL efficacy Granuloma-scale, Early \| Late |
|---|---|---|---|---|---|---|---|---|---|---|---|
| Fraction of macrophage-released bacteria that get trapped in caseum (CN) | | | - | | | | | | | | - |
| Growth rate constant of extracellular bacteria (alpha20) | - | | | | | | | | - | | - |

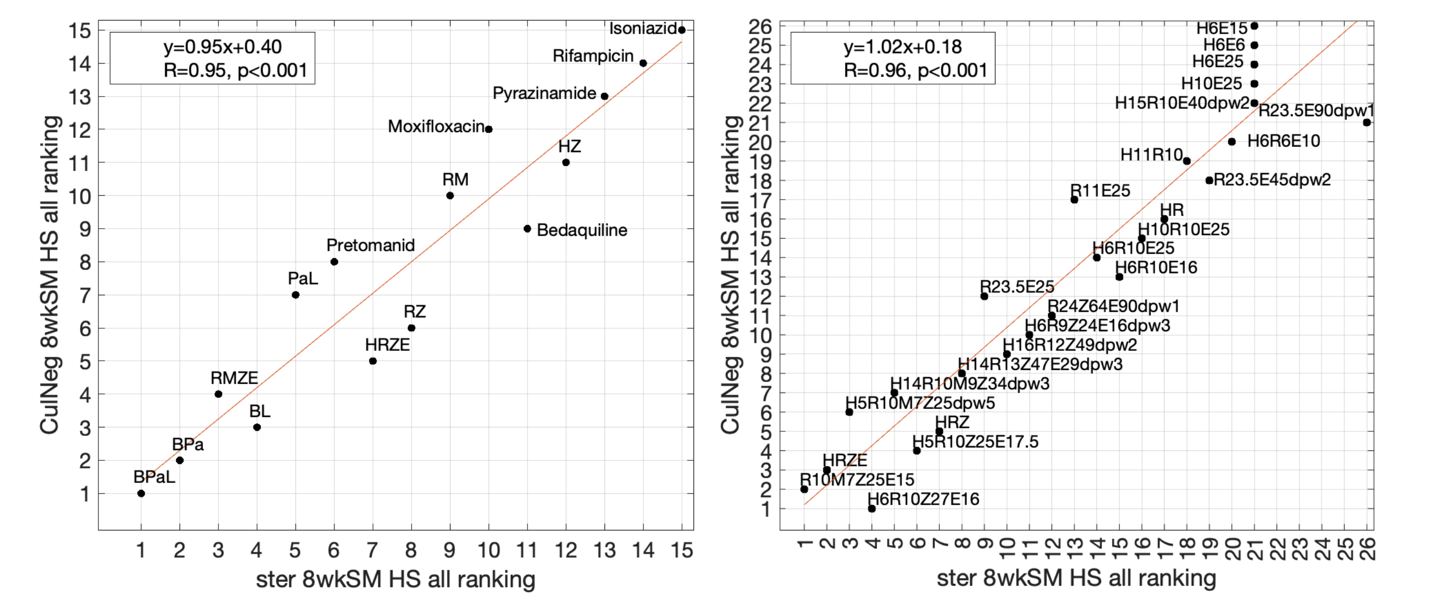

**Supplemental Figure 1**: Host-scale sterilizing and culture-negative rankings of regimen sets (ii) and (iii) are strongly correlated.
